# A neural circuit linking two sugar sensors regulates satiety-dependent fructose drive in *Drosophila*

**DOI:** 10.1101/2021.04.08.439043

**Authors:** Pierre-Yves Musso, Pierre Junca, Michael D Gordon

## Abstract

Ingestion of certain sugars leads to activation of fructose sensors within the brain of flies, which then sustain or terminate feeding behavior depending on internal state. Here, we describe a three-part neural circuit that links satiety with fructose sensing. We show that AB-FBl8 neurons of the Fan-shaped body display oscillatory calcium activity when hemolymph glycemia is high, and that these oscillations require synaptic input from SLP-AB neurons projecting from the protocerebrum to the asymmetric body. Suppression of activity in this circuit, either by starvation or genetic silencing, promotes specific drive for fructose ingestion. Moreover, neuropeptidergic signaling by tachykinin bridges fan-shaped body activity and Gr43a-mediated fructose sensing. Together, our results demonstrate how a three-layer neural circuit links the detection of two sugars to impart precise satiety-dependent control over feeding behavior.

## INTRODUCTION

Animals have evolved multi-layered systems to ensure fulfillment of their nutrient requirements. Among necessary nutrients, carbohydrates afford essential energy and play a major role in survival. Circulating glucose provides fast available energy to support tissue function, and glycemia is tightly regulated. Small variations impact food drive, while prolonged deficiencies lead to tissue damage. Glycemic homeostasis is a well-studied phenomenon under hormonal control, but ultimately the only way to gain energy is through feeding.

Food seeking in response to lowered glycemia is a behavioral consequence that is conserved between mammals and *Drosophila* (Dus et al., 2013; 2011; Niwano et al., 2009). In flies, several neurons have been suggested to directly sense glucose and impact feeding behavior, including those expressing the peptide DH44 (Dus et al., 2015) and a set co-expressing peptides Corazonin (Crz) and short neuropeptide F (sNPF) dubbed “*CN* neurons” (Kapan et al., 2012; Oh et al., 2019). Both populations display oscillatory calcium activity in the presence of circulating nutritive sugars and are necessary for post-ingestive nutrient selection (Dus et al., 2015; Oh et al., 2019). MB-MP1 neurons of the mushroom bodies (MBs) also display calcium oscillations, which are thought to signal the availability of energy required for long-term memory formation (Musso et al., 2015; Plaçais and Preat, 2013; Plaçais et al., 2012).

How do changes in circulating glucose alter feeding? The most straightforward way is to elicit changes in sensitivity and attractiveness to specific food-related cues. Important examples of this phenomenon have been described in both the olfactory system (Ko et al., 2015; Lin et al., 2019; Root et al., 2011; Sayin et al., 2019; Slankster et al., 2020), and the gustatory system of flies (Inagaki et al., 2012; 2014; Jaeger et al., 2018; LeDue et al., 2016; Musso et al., 2019). In taste, a particular area of focus has been sugar detection. Sugars are first detected by gustatory receptor neurons located on the fly proboscis and legs, which initiate feeding, and then by GRNs in the pharynx, which sustain ingestion (Dahanukar et al., 2001; 2007; Jiao et al., 2008; Kwon et al., 2014; LeDue et al., 2015; Marella et al., 2006; Park and Kwon, 2011; Scott, 2018; Thorne et al., 2004; Wang et al., 2004b). The responses of sugar-sensing GRNs and several examples of second-order sugar neurons are enhanced by food deprivation (Inagaki et al., 2014; Kain and Dahanukar, 2015; Yapici et al., 2016).

Starvation also modulates the response of neurons in the lateral protocerebrum of the brain that express one particular member of the gustatory receptor (GR) family, Gr43a (Miyamoto et al., 2012). Gr43a is unusual among the nine identified sugar-sensing GRs in that it is specifically tuned to fructose. Activation of Gr43a brain neurons leads to feeding cessation in fed flies and feeding prolongment in those that are starved (Miyamoto et al., 2012). This supports a model where circulating fructose levels rise when a fly eats sugary food, and trigger activation of Gr43a brain neurons that then either prolong or terminate feeding, depending on satiety state. One appealing aspect of this model is that it cleanly separates detection of ingested sugars from detection of satiety. Naturally-occurring sweet foods generally contain both fructose and glucose, as well as sucrose, which is a dimer of the two. Thus, circulating fructose may serve as the cue for recently ingested sugar and vary widely, while tightly-controlled glucose levels provide an indicator of satiety. However, the circuitry connecting starvation and glycemia with fructose sensing by Gr43a brain neurons has not been explored.

In this study, we find that AB-FBl8 (vΔA_a) neurons, which compose layers 8 and 9 of the central complex structure called the Fan-Shaped body, act as central glucose sensors that couple satiety state with fructose drive. Fed flies display AB-FBl8 calcium oscillations and have equal preference between feeding on fructose or glucose. However, prolonged starvation suppresses AB-FBl8 oscillations and leads to a strong shift in preference towards fructose. We show that silencing AB-FBl8 neurons mimics the fructose preference shift seen upon prolonged starvation and that AB-FBl8 activity is modulated by glutamatergic input from SLP-AB neurons of superior lateral protocerebrum. Finally, we demonstrate that the effect of AB-FBl8 neurons on fructose feeding is mediated by release of the neuropeptide tachykinin, which signals to Gr43a brain neurons. The linking of two specific sugar sensors in this three-neuron circuit imparts precise hunger-dependent control over sugar consumption.

## RESULTS

### Starvation regulates Fan-Shaped Body oscillations

In order to identify novel circuits controlling carbohydrate intake, we surveyed calcium activity in candidate brain areas using GCaMP6f expression under control of GAL4 lines from the Janelia Flylight collection (Chen et al., 2013; Jenett et al., 2012). This revealed oscillatory activity in dorsal fan-shaped body (dFB) neurons labelled by *R70H05-GAL4* (Figure 1A). Oscillations were strong in fed flies and dramatically reduced in intensity and frequency following prolonged starvation of 30 hours (Figure 1B-F; Figure S1A,B; supplemental movies 1,2). We noted asymmetry in the oscillations, with asynchronous activity on the right and left side as well as a tendency for the right part of the dFB to show higher frequencies than the left (Figure S1A,B; supplemental movie 1). Interestingly, flies starved for only 18 hours displayed oscillations similar to those of fed flies (Figure 1B-F; Figure S1A,B). Since starvation is associated with lower hemolymph carbohydrate levels (Dus et al., 2011; 2013; Rovenko et al., 2015a; 2015b), we hypothesized that dFB neurons may act as brain glucose sensors. Knocking down Glucose transporter type 1 (Glut1) and Hexokinase C (HexC) within dFB reduced their oscillations to a level comparable to prolonged starvation, supporting a role for glucose sensing in dFB regulation (Figure 1B-G; Figure S1A,B) (Dus et al., 2015; Escher and Rasmuson-Lestander, 1999; Moser et al., 1980; Oh et al., 2019; Volkenhoff et al., 2018).

**Figure 1.**
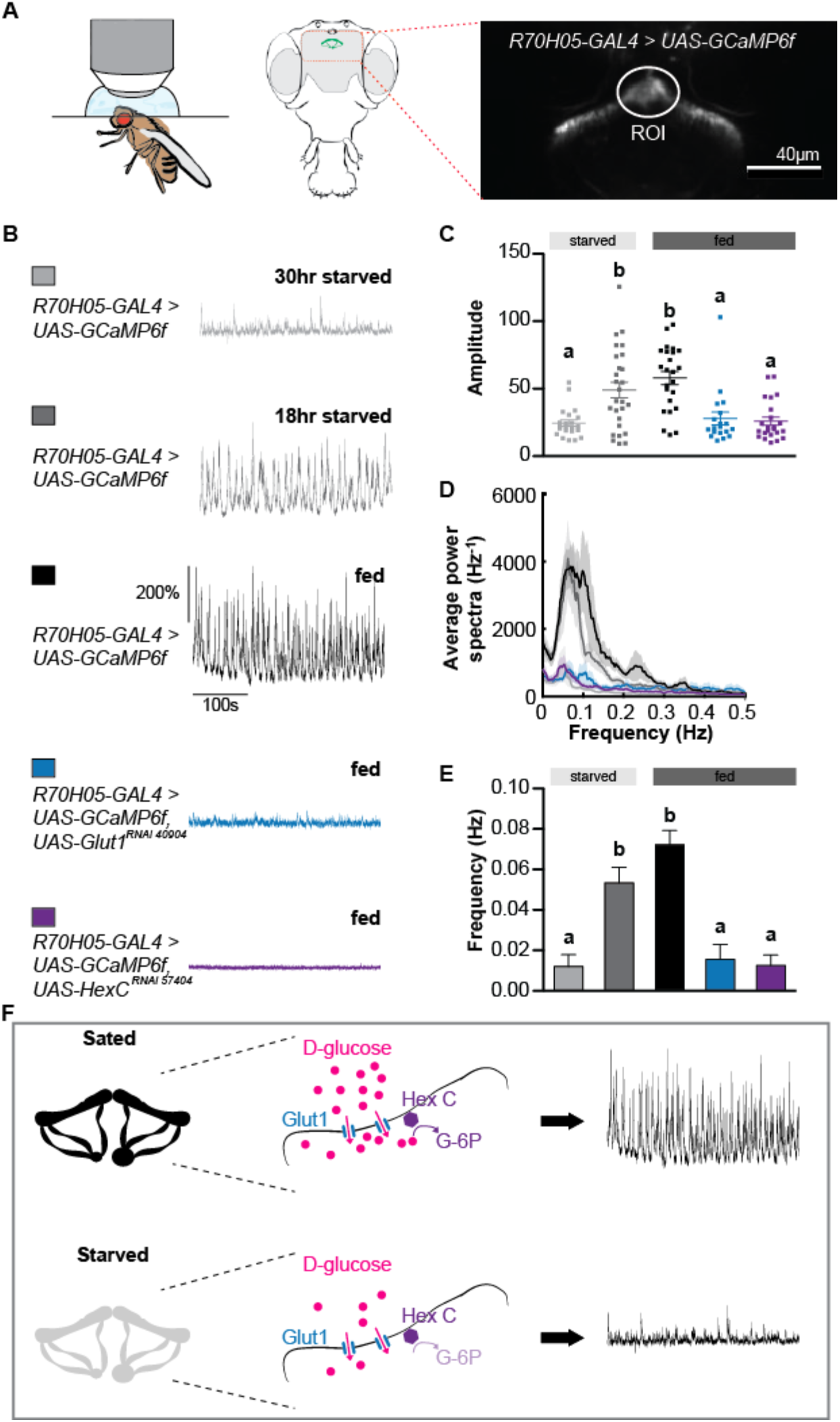
Starvation regulates Fan-Shaped Body oscillations. (A) Schematic of imaging preparation to monitor calcium oscillations (left) with GCaMP6f signal from *R70H05-GAL4* expression in the dFB (right). (B) Calcium traces from *R70H05-GAL4 > UAS-GCamP6f* flies after different periods of starvation or expressing RNAi against Glut1 and HexC. (C) Amplitudes of dFB neurons oscillations. (D) Power spectra of dFB neurons oscillations. (E) Frequencies of dFB neurons oscillations. (F) Model: in sated flies, D-glucose enter the dFB neurons through Glut1 and trigger oscillations through the activity of HexC; in starved flies, the low availability of D-glucose prevents oscillations. Values represent mean ± SEM. *n* = 19-27. Statistical tests: one-way ANOVA and Tukey post-hoc; different letters represent significant differences *p* < 0.05. See also Figure S1.

### AB-FBl8 drives starvation-dependent changes in fructose feeding

The dFB neurons labelled by *R70H05-GAL4* occupy layers 8 and 9 of the FB and project to the Asymmetric Body (AB), identifying them as AB-FBl8 (or vΔA_a) neurons (Figure 1A) (Hulse et al., 2020; Jenett et al., 2012; Wolff and Rubin, 2018; Wolff et al., 2015). To test whether AB-FBl8 neurons link satiety signals to changes in behavior, we silenced them and measured feeding using a modified version of FlyPAD where food interactions were calculated using the algorithm we developed for the STROBE (Figure 2B-C) (Itskov et al., 2014; Jaeger et al., 2018; Musso et al., 2019). Flies conditionally expressing the inward rectifying potassium channel Kir2.1 (Baines et al., 2001; McGuire et al., 2004) under the control of *R70H05-GAL4* displayed an increased number of interactions with sucrose at low concentrations (5 and 50 mM) but not at 1 M (Figure 2D). Interestingly, AB-FBl8 silencing did not affect feeding on either L-glucose (sweet but not caloric) or sorbitol (caloric but not sweet), but dramatically elevated interactions with a mixture of the two (Figure 2E-G) (Dus et al., 2013; Musso et al., 2015; 2017). Given that the sweet taste of sucrose and L-glucose stimulates feeding initiation, we suspected that sorbitol and sucrose were being detected post-ingestively to trigger enhanced feeding in AB-FBl8-silenced flies. D-glucose failed to trigger excess feeding by AB-FBl8-silenced flies, indicating that energy alone was not sufficient for this post-ingestive effect (Figure 2H). However, AB-FBl8 silencing produced strong over feeding on fructose, which is quickly metabolized from both sucrose and sorbitol (Figure 2I). Thus, we posited that AB-FBl8 are part of a circuit that links satiety-dependent changes in hemolymph glucose levels with a flies’ response to post-ingestive changes in hemolymph fructose.

**Figure 2.**
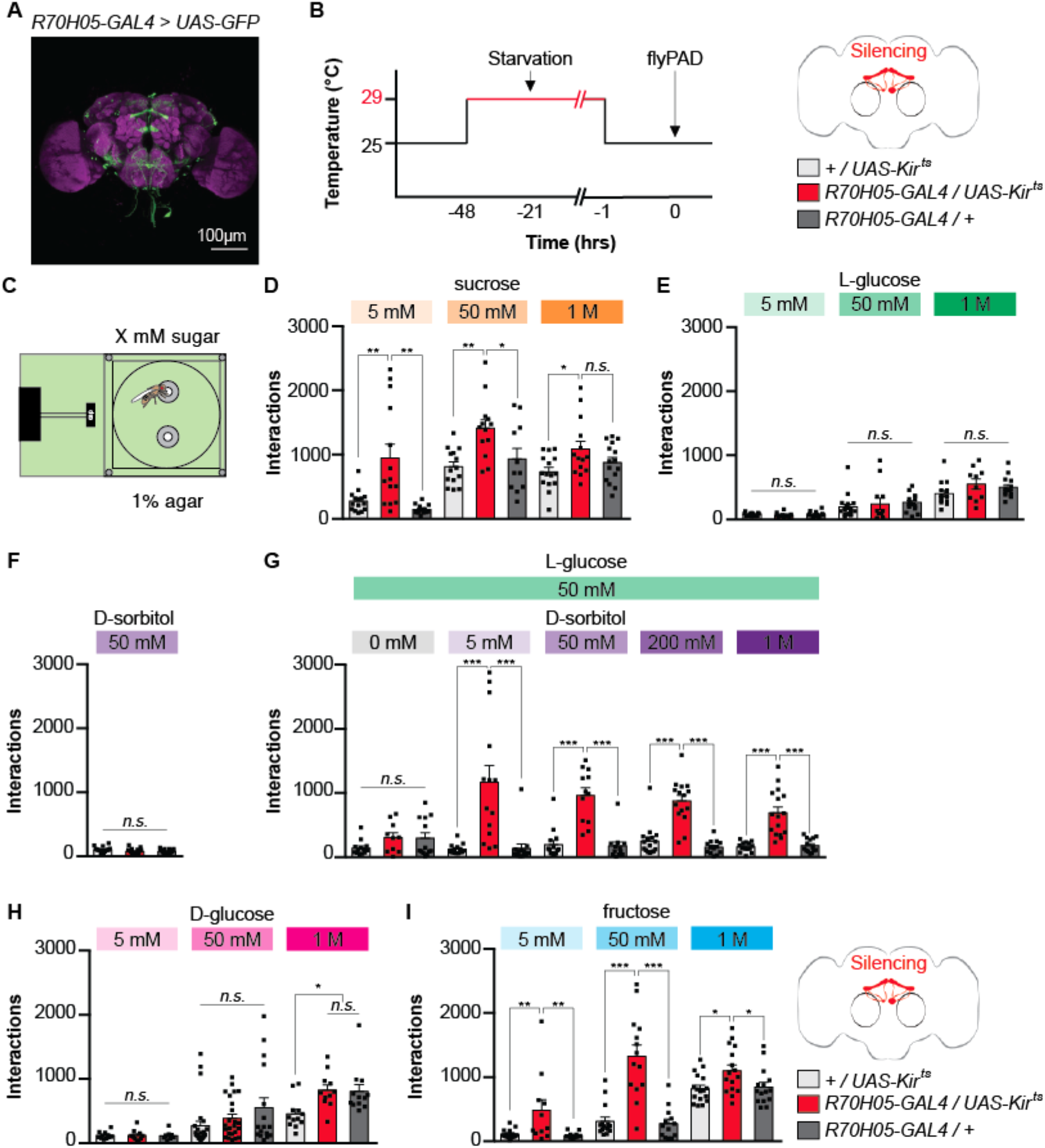
Silencing AB-FBl8 neurons increases fructose feeding. (A) Immunofluorescent detection of *UAS-GFP* driven by *R70H05-GAL4*. (B) Experimental timeline: flies are placed at 29 degrees for 47 hrs and starved for 18 hrs, and experiments are performed at 25 degrees. (C) Experimental setup: one channel is filled with sugar and the other one is filled with 1% agar. (D) Effect of silencing AB-FBl8 neurons on flies’ interactions with various concentrations of sucrose (5, 50 and 1000 mM; *n* = 16-21). (E) Effect of silencing AB-FBl8 neurons on flies’ interactions with various concentrations of L-glucose (5, 50 and 1000 mM; *n* = 10-19). (F) Effect of silencing AB-FBl8 neurons on flies’ interactions with 50 mM of D-sorbitol (*n* = 15). (G) Effect of silencing AB-FBl8 neurons on flies’ interactions with 50 mM of L-glucose mixed with various concentrations of D-sorbitol (0, 5, 50, 200 and 1000 mM; *n* = 10-16). (H) Effect of silencing AB-FBl8 neurons on flies’ interactions with various concentrations of D-glucose (5, 50 and 1000 mM; *n* = 8-26). (I) Effect of silencing AB-FBl8 neurons on flies’ interactions with various concentrations of fructose (5, 50 and 1000 mM; *n* = 11-17). Values represent mean ± SEM. Statistical tests: one-way ANOVA and Tukey post-hoc; ns: *p* > 0.05, \**p* < 0.05, ** *p* < 0.01, \*\*\**p* < 0.001.

If AB-FBl8 produces satiety signals that inhibit post-ingestive fructose sensing, then silencing of AB-FBl8 should release this inhibition and produce increased relative preference for fructose over glucose. We found that flies expressing Kir2.1 under control of *R70H05-GAL4* displayed a strong preference for fructose over glucose at concentrations of 5 mM and 50 mM, but not 1 M, while control flies showed nearly equal preference for the two sugars at all concentrations (Figure 3A-B; Figure S1C). Since *R70H05-GAL4* drives expression in additional neurons outside the AB-FBl8 population, we verified the causal role for AB-FBl8 by measuring glucose versus fructose preference following silencing with two other drivers (*R70H05-LexA* and *VT005528-GAL4*) and a split construction that we built (*FB-split*), all of which specifically label AB-FBl8 neurons (Figure S1D-F). In each case, AB-FBl8 silencing led to a strong preference for fructose over glucose. We also verified that AB-FBl8 neurons labeled by *FB-split* showed oscillatory activity that was reduced following prolonged starvation (Figure S2A-C).

**Figure 3:**
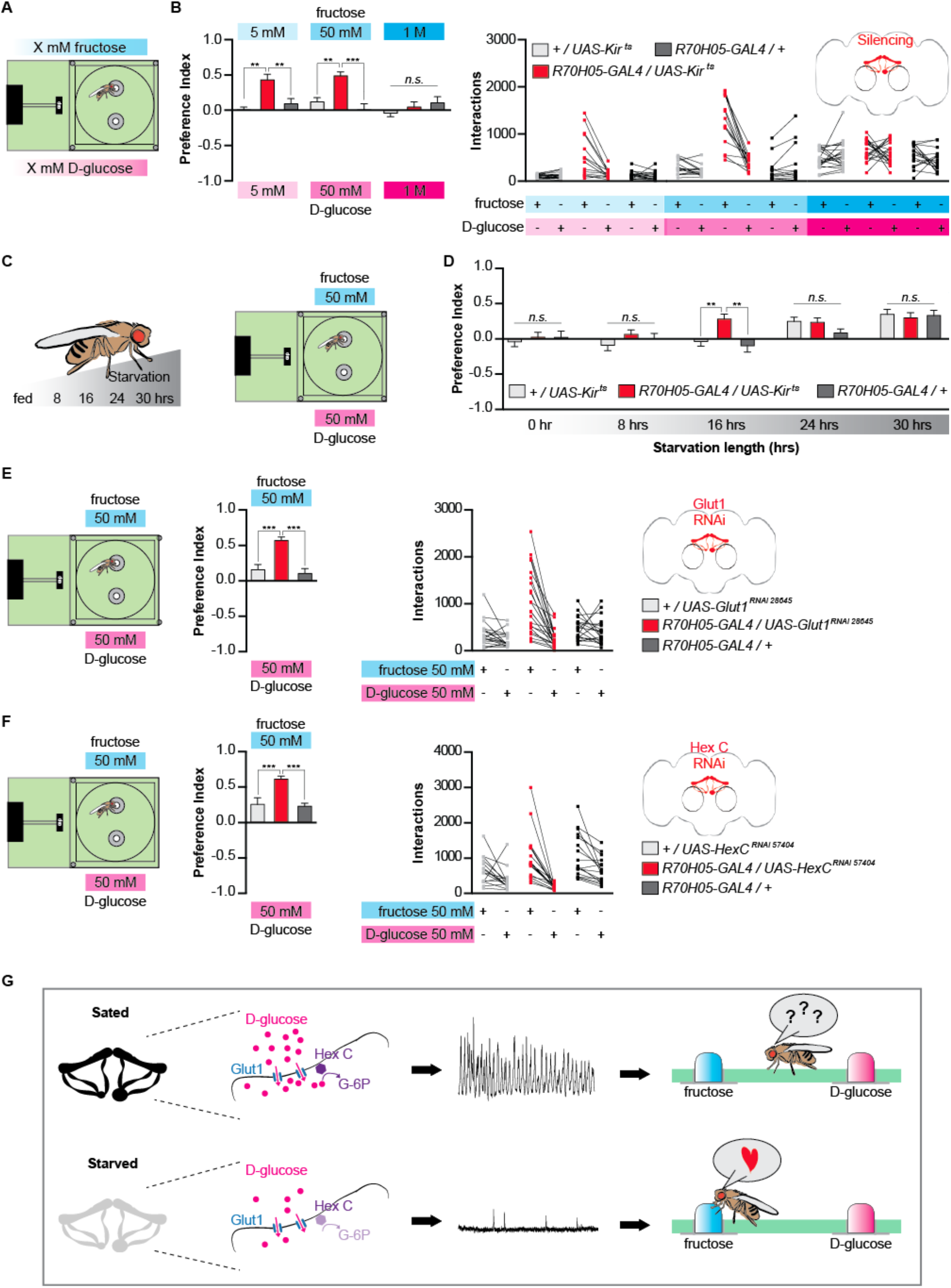
Fructose feeding preference relies on starvation. (A) Experimental setup: left channel is filled with fructose and the right channel is filled with the same concentration of D-glucose. (B) Effect of silencing AB-FBl8 neurons on flies’ preference between fructose and D-glucose at 5 mM, 50 mM, and 1 M (left); and their corresponding interactions (right; *n* = 13-21). (C) Experimental setup: Flies are starved for different amounts of time and given the choice between 50 mM fructose and 50 mM D-glucose. (D) Effect of starvation length on preference between 50 mM fructose and 50 mM D-glucose in controls and flies with silenced AB-FBl8 neurons (*n* = 21-36). (E) Effect of knocking down Glut1 in AB-FBl8 neurons on the preference between 50 mM fructose and D-glucose (*n* = 19-23). F) Effect of knocking down HexC in AB-FBl8 neurons on the preference between 50 mM fructose and D-glucose (*n* = 15-18). (G) Model for regulation of fructose preference by AB-FBl8 activity. Values represent mean ± SEM. Statistical tests: *one-way ANOVA* and *Tukey post-hoc*; ns: *p* > 0.05, ** *p* < 0.01, \*\*\**p* < 0.001. See also Figure S1, S2 and S3.

Given that AB-FBl8 oscillations are suppressed upon starvation, we wondered whether AB-FBl8 silencing evoked a starvation-like state in flies. Thus, we subjected AB-FBl8-silenced and control flies to different periods of food deprivation and then measured their preference for fructose versus glucose (Figure 3C). This revealed that both AB-FBl8-silenced and control flies prefer fructose over glucose following food deprivation of 24 hours or more (Figure 3D). However, at 16 hours of starvation, a condition under which AB-FBl8 show fed-like oscillations (Figure 1B), AB-FBl8-silenced flies preferred fructose, while control flies fed on both options equally. These results indicate that flies develop a relative preference for fructose as starvation progresses, and that AB-FBl8 silencing shifts this curve to the left, producing fructose preference at lower levels of starvation (Figure 3D). Notably, preference between fructose and glucose was equal for all groups following 0 or 8 hours of starvation. We suspect that this is because a threshold of consumption needs to be met for post-ingestive fructose sensing to stimulate further feeding, and that feeding initiation is controlled independently of AB-FBl8 activity. Thus, flies without sufficient food deprivation would not consume enough fructose to trigger AB-FBl8-regulated feeding circuits. Indeed, the proboscis extension reflex (PER) to fructose and glucose remained unchanged following AB-FBl8 silencing, indicating that AB-FBl8 do not regulate peripheral sensitivity to sugars or sensory-driven feeding initiation, and likely rather modulate responses to post-ingestive cues (Figure S2D,E).

Although food interactions measured on the FlyPAD strongly correlate with consumption, we next sought to confirm that AB-FBl8 silencing genuinely promotes fructose ingestion. As expected, AB-FBl8-silenced flies preferentially consumed fructose over glucose in a dye-based binary choice feeding assay, while controls consumed the two sugars equally (Figure S2F). Moreover, control flies preferentially consumed fructose over glucose when AB-FBl8 activity was suppressed by 43 hours of starvation, and thermogenetic activation of AB-FBl8 with TRPA1 suppressed this elevated fructose feeding (Figure S2G). This demonstrates that AB-FBl8 activity is sufficient to inhibit fructose sensing mechanisms.

To link the behavioral role of AB-FBl8 back to their function in glucose sensing, we measured the feeding preference of flies following RNAi knock down of Glut1 or HexC in the AB-FBl8. Consistent with their effects on AB-FBl8 oscillations, knock-down of either gene promoted strong preference for fructose over glucose in the FlyPAD (Figure 3E-F; Figure S3A-B). This suggests that changes in AB-FBl8 activity mediated by glucose sensing drive effects on fructose feeding. Altogether, our results suggest that starvation-induced reduction of hemolymph glucose suppresses AB-FBl8 activity, which in turn promotes fructose feeding (Figure 3G).

### SLP-AB synaptically modulate AB-FBl8 oscillations

To examine the broader circuit in which AB-FBl8 are regulating fructose consumption, we used *UAS-synaptotagmin-GFP* (*UAS-Syt-GFP*) and *UAS-Denmark* to label pre- and post-synaptic areas, respectively. This demonstrated that the AB-FBl8 presynaptic terminals reside in layer 8 of the Fan-Shaped Body, while their dendrites primarily occupy the AB, a structure known to be required for energetically-costly long-term memory (Figure 4A)(Burke and Waddell, 2011; Mery and Kawecki, 2005; Musso et al., 2015; Pascual et al., 2004; Plaçais and Preat, 2013; Plaçais et al., 2017). In search of inputs to AB-FBl8, we examined the SLP-AB population, which has arborizations in the Superior lateral protocerebrum (SLP) and the AB (Jenett et al., 2012; Wolff and Rubin, 2018). We generated a split-GAL4 labelling SLP-AB neurons, and confirmed the location of their dendrites in the SLP and axon terminals in the AB (Figure 4B). Trans-Tango driven by this driver revealed post-synaptic neurons in layer 8 of the Fan-Shaped Body, suggesting that AB-FBl8 are postsynaptic to SLP-AB (Figure 4C)(Talay et al., 2017). Moreover, GFP-reconstitution across synaptic partners (GRASP) between AB-FBl8 and SLP-AB revealed a single point of contact in the AB (Figure 4D). Finally, silencing SLP-AB neurons reduced the oscillatory activity of AB-FBl8 in fed flies, demonstrating functional connectivity between the two neuron populations (Figure 4E,H; Figure S4).

**Figure 4.**
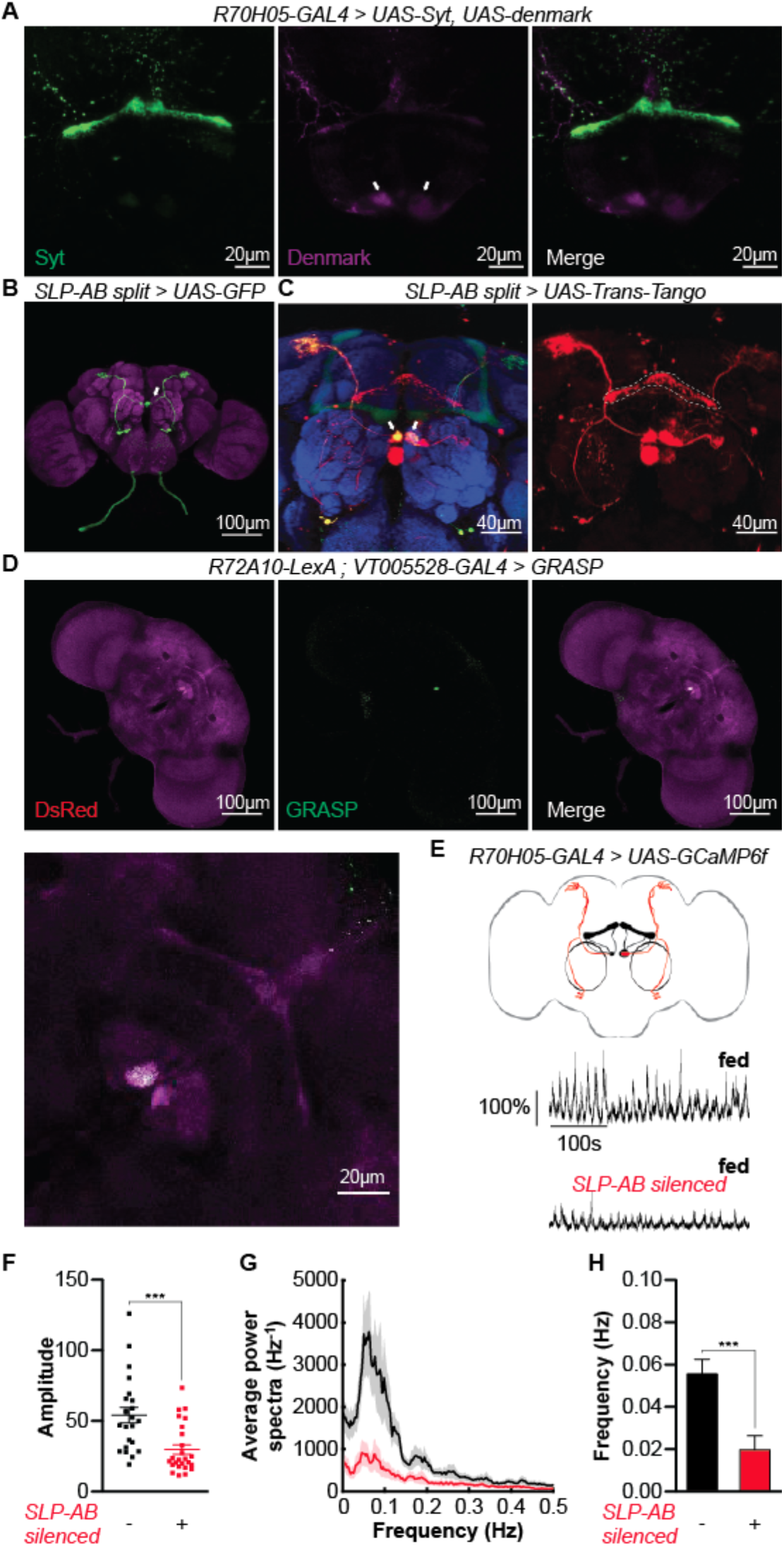
SLP-AB neurons contact AB-FBl8 neurons on the asymmetric body and modulate oscillations. (A) Immunofluorescent detection of *UAS-Syt* (green) and *UAS-Denmark* (magenta) driven by *R70H05-GAL4*. Arrows show dendritic compartment localized on the asymmetric body. (B) Immunofluorescent detection of *UAS-GFP* driven by *SLP-AB* split-GAL4. Arrows show projections to the asymmetric body. (C) Trans-Tango expression driven by SLP-AB split-Gal4. Arrows show the contact between SLP-AB neurons and post synaptic targets, and dotted line outlines trans-Tango expression in the Fan-shaped body layer 8. (D) GRASP between AB-FBl8 and SLP-AB neurons produces a signal at the asymmetric body. (E) Calcium trace from *R70H05-GAL4 > UAS-GCamP6f* (top) and *R70H05-GAL4; R72A10-LexA > UAS-GCamP6f; LexAop-tnt* (bottom) fed flies. (F) Amplitudes of oscillations. (G) Power spectra of oscillations. (H) Frequencies of oscillations. Imaging data are from ROI placed in the middle part of AB-FBl8 neurons. Values represent mean ± SEM. *n* = 23-25. Statistical tests: *t*-test, \*\*\**p* < 0.001. See also Figure S4.

Next, we addressed whether the modulatory action of SLP-AB neurons on AB-FBl8 affects behavior. Indeed, silencing SLP-AB neurons reproduced all the behavioral phenotypes observed from AB-FBl8 silencing with *R70H05-GAL4*: increased feeding interactions with sucrose, fructose, and a mixture of L-glucose and sorbitol, but not D-glucose, L-glucose, or sorbitol alone (Figure S5A-H); no effect on taste sensitivity to fructose nor D-glucose (Figure S5I-J); and enhanced preference for fructose over glucose (Figure 5A; Figure S5K). Silencing SLP-AB with an independent driver (*R72A10-LexA* (Jenett et al., 2012)) also reproduced the fructose feeding preference, verifying that SLP-AB were responsible for this phenotype (Figure S5L). Moreover, like AB-FBl8, SLP-AB activation reduced fructose feeding preference in strongly starved flies (Figure S5M).

**Figure 5.**
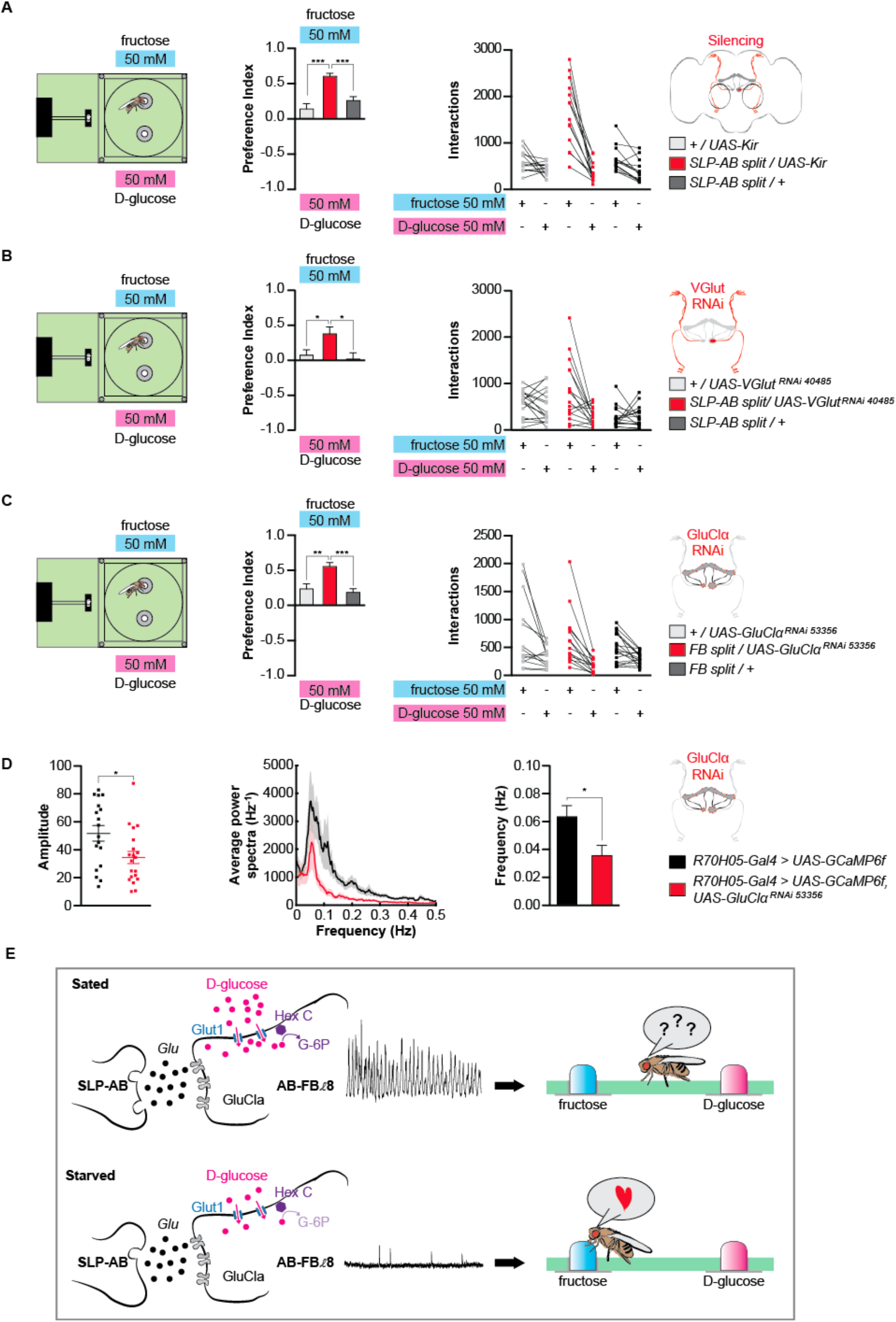
Glutamatergic SLP-AB neurons are positively connected to AB-FBl8 neurons. (A) Effect of silencing SLP-AB neurons on flies’ preference between fructose and D-glucose, and their corresponding interactions (*n* = 13-14). (B) Effect of knocking down *Vglut* in SLP-AB neurons on flies’ preference between fructose and D-glucose, and their corresponding interactions (*n* = 16-19). (C) Effect of knocking down *GluClα* in AB-FBl8 neurons on flies’ preference between fructose and D-glucose, and their corresponding interactions (*n* = 14-19).(D) Knocking down *GluClα* in AB-FBl8 neurons moderately inhibits oscillations (*n* = 18-20). (E) Glutamatergic input from SLP-AB is permissive for AB-FBl8 oscillations). Values represent mean ± SEM. Statistical tests: one-way ANOVA and Tukey post-hoc for behavior and *t*-test for imaging; ns: *p* > 0.05, ** *p* < 0.01, \*\*\**p* < 0.001. See also Figure S5 and S6.

Knocking down Glut1 and Hexokinase C in SLP-AB did not impact behavior, suggesting that these neurons do not sense hemolymph D-glucose (Figure S6A,B). However, knocking down the vesicular glutamate transporter (Vglut) in SLP-AB promoted fructose feeding preference (Figure 5B; Figure S6C-E). GluClα, a ligand gated chloride channel, is one of the few glutamate receptors expressed in FB (Cully et al., 1996; Kahsai et al., 2012; Kondo et al., 2020). Knock-down of GluClα in AB-FBl8 also induced fructose preference and decreased AB-FBl8 oscillations (Figure 5C,D; Figure S6F-H). Altogether, these data suggest that SLP-AB impact behavior by promoting AB-FBl8 oscillations via the action of glutamate on GluClα(Figure 5E).

### AB-FBl8 regulates fructose feeding via tachykinin signalling to Gr43a neurons

Initiating trans-tango from the AB-FBl8 did not show any clear post-synaptic targets, with the exception of the noduli (Figure S7A,B). Such an absence of trans-tango signal suggests that AB-FBl8 may function non-synaptically through peptide secretion, which is characteristic of calcium oscillatory cells (Thorner et al., 1988). This is supported by EM data revealing postsynaptic connections only within the FB and the AB, and a previous report of tachykinin and sNPF expression in AB-FBl8 (Clements et al., 2020; Kahsai et al., 2012; Kahsai and Winther, 2011; Qi et al., 2020; Winther et al., 2003). Using two independent AB-FBl8 drivers, we found that knock down of Tk, but not sNPF, reproduced the fructose preference phenotype seen with AB-FBl8 silencing (Figure 6 and Figure S7C,D). Moreover, a recent study showed elevated Tk mRNA expression in fed and re-fed flies compared to those that had been starved (Qi et al., 2020).

**Figure 6.**
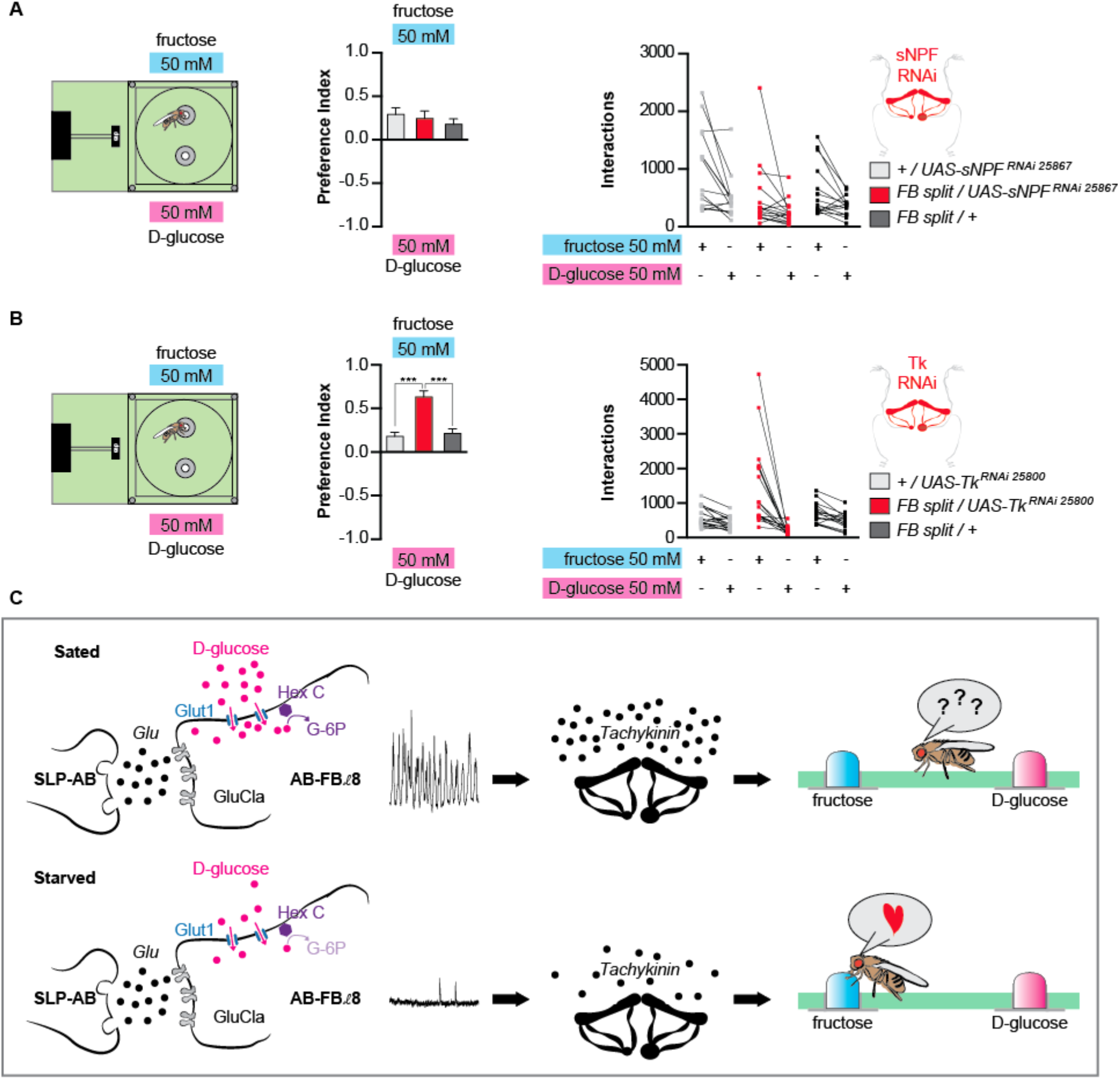
AB-FBl8 neurons regulate fructose-feeding preference through tachykinin secretion. (A) Effect of knocking down sNPF in AB-FBl8 neurons on flies’ preference between fructose and D-glucose, and their corresponding interactions (*n* = 14-18). (B) Effect of knocking down tachykinin in AB-FBl8 neurons on flies’ preference between fructose and D-glucose, and their corresponding interactions (*n* = 16-20). (C) Tachykinin secretion in sated flies inhibit fructose feeding preference. Values represent mean ± SEM. Statistical tests: one-way ANOVA and Tukey post-hoc; ns: *p* > 0.05, \*\*\**p* < 0.001. See also Figure S7.

Flies express two receptors for Tk: TkR86C (or NKD)(Asahina et al., 2014; Li et al., 1991; Monnier et al., 1992; Poels et al., 2009) and the widely expressed TkR99D (or DTKR) (Birse et al., 2006; Ignell et al., 2009; Im et al., 2015; Qi et al., 2020; Song et al., 2014). Since Gr43a-expressing neurons in the lateral protocerebrum are the only known post-ingestive fructose sensors, we knocked down each receptor specifically in Gr43 brain neurons using *Gr43a-GAL4* combined with Cha^7.4kb^-GAL80 (Miyamoto et al., 2012) (Figure S8A-B). This revealed a requirement for TkR99D but not TkR86C in restricting fructose intake (Figure 7A,C and Figure S8). In vivo, TkR99D has been demonstrated to have inhibitory activity (Birse et al., 2006; Ignell et al., 2009; Song et al., 2014). Thus, we postulate that under fed conditions, Tk released from AB-FBl8 inhibits brain Gr43a neurons, preventing them from responding to circulating fructose (Figure 7C).

**Figure 7.**
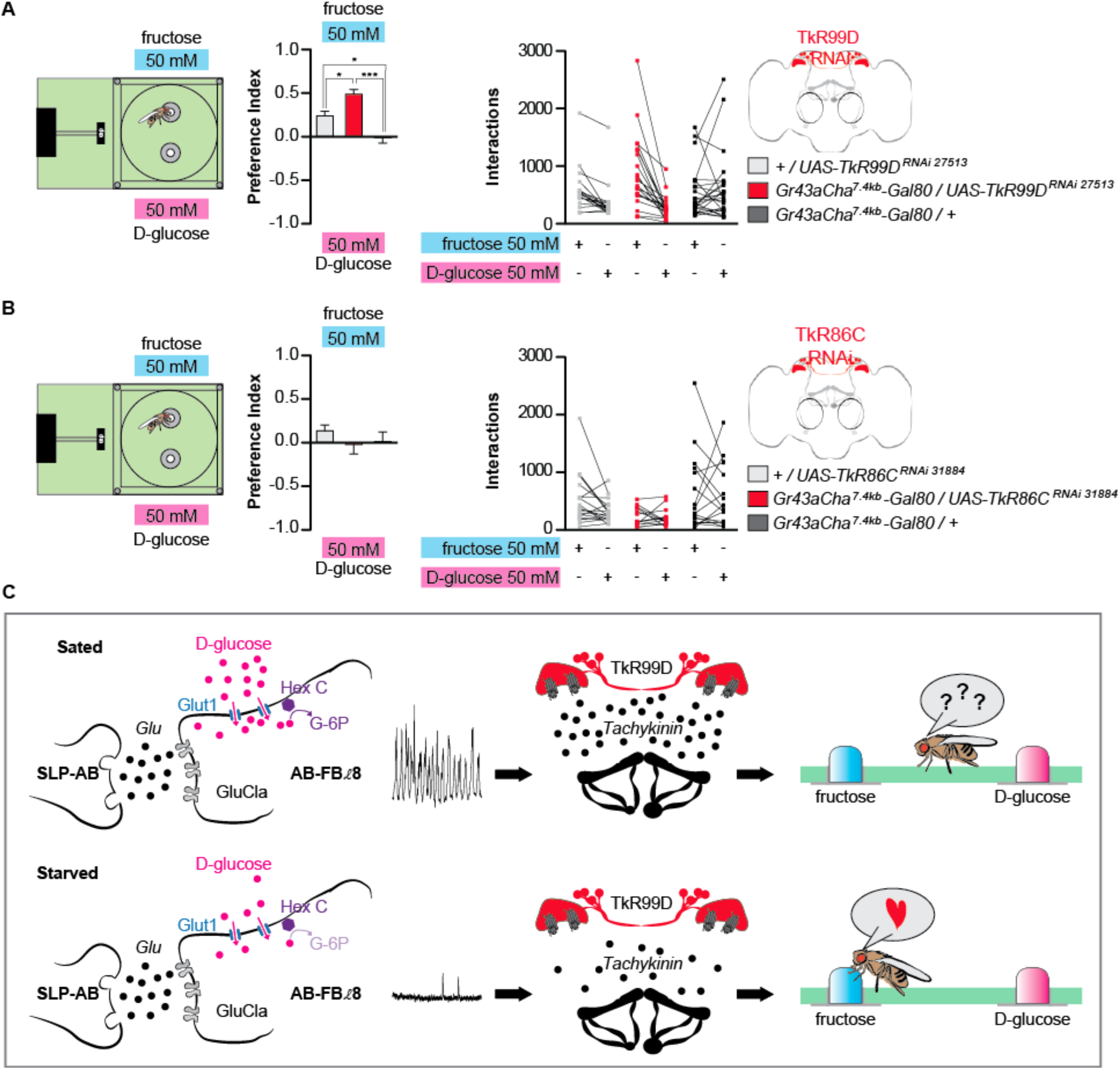
AB-FBl8 neurons’ tachykinin secretion regulates fructose feeding preference through activation of the brain fructose sensor Gr43a. (A) Effect of knocking down TkR99D in brain Gr43a neurons on flies’ preference between fructose and D-glucose, and their corresponding interactions (*n* = 18-26). (B) Effect of knocking down TkR86C in brain Gr43a neurons on flies’ preference between fructose and D-glucose, and their corresponding interactions (*n* =14-19). (C) Model: tachykinin secretion in sated flies inhibits fructose feeding preference by acting on TkR99D expressed in Gr43a neurons. Values represent mean ± SEM. Statistical tests: one-way ANOVA and Tukey post-hoc; ns: *p* > 0.05, \**p* < 0.05, \*\*\**p* < 0.001. See also Figure S8.

### Brain Gr43a neurons acutely regulate fructose feeding

To understand how Gr43a brain neurons impact feeding, we first tested their role in the choice between fructose and glucose using the FlyPAD. Flies expressing Kir2.1 under control of *Gr43a-GAL4* and *Cha^7.4kb^-GAL80* strongly prefer glucose over fructose, indicating that Gr43a brain neurons are necessary for normal fructose feeding (Figure 8A). Conversely, closed-loop activation of Gr43a brain neurons in the sip-triggered optogenetic behavior enclosure (STROBE) is sufficient to promote feeding (Figure 8B). In this experiment, flies expressing the red-light activated channel CsChrimson in Gr43a brain neurons and previously fed the obligate CsChrimson cofactor all-trans retinal were compared to a control group without retinal (Klapoetke et al., 2014). Flies could feed on either of two identical drops of 1% agar, one of which was coupled to red light activation. We found that the retinal-fed group robustly preferred the light-triggering agar, while the control group showed no preference (Figure 8B).

**Figure 8.**
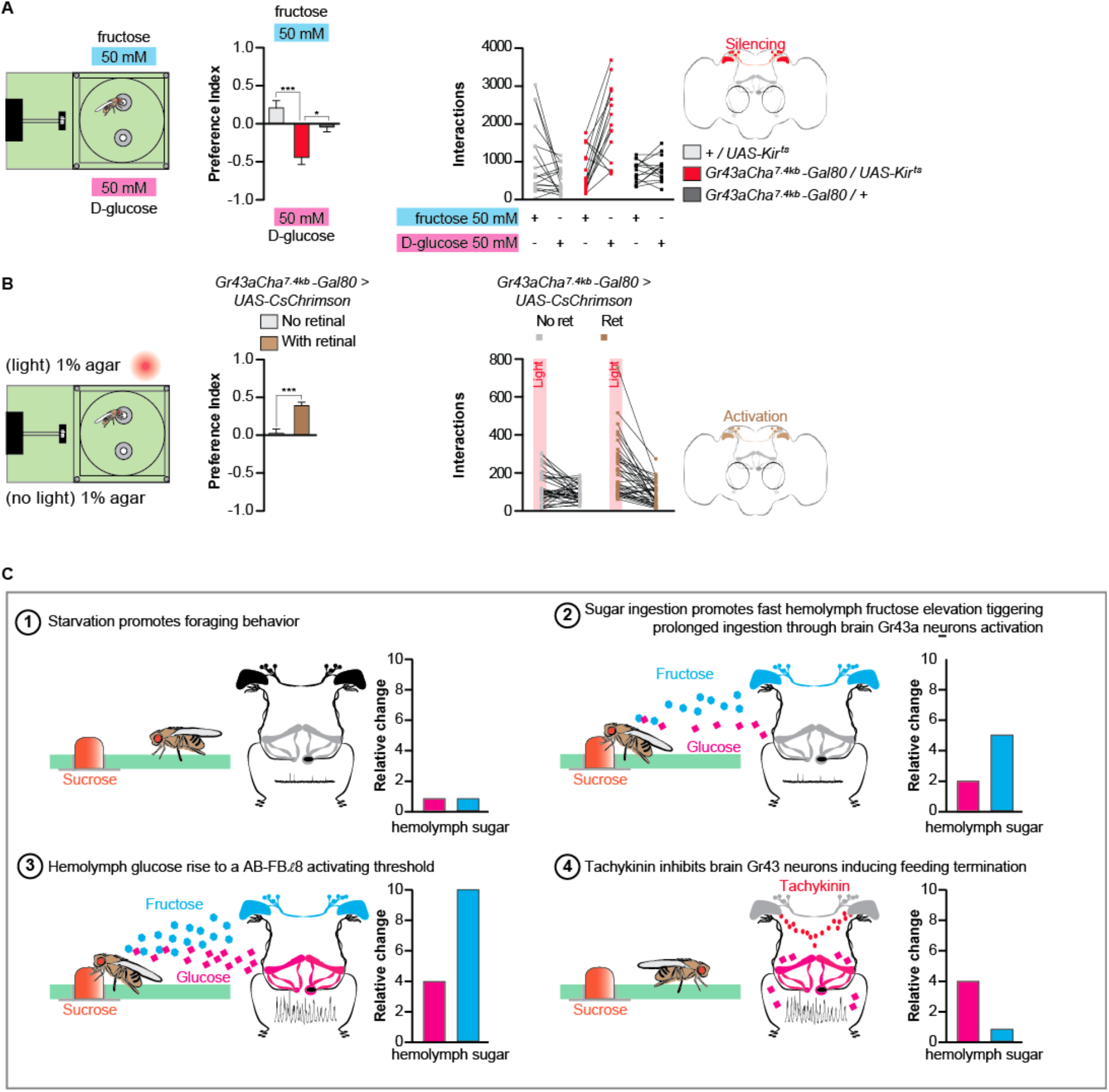
Brain Gr43a neurons acutely regulate fructose feeding. (A) Effect of silencing brain Gr43a neurons on flies’ preference between fructose and D-glucose, and their corresponding interactions (*n* = 15-19). (B) Preference in the STROBE for agar paired with light activation of Gr43a neurons expressing CsChrimson, and their corresponding interactions (*n* = 40-41). (C) Model for how fructose serves as a cue for promoting sugar ingestion, and how rising glucose levels signal satiety through AB-FBl8, which then inhibits sensitivity to fructose and terminates feeding. Values represent mean ± SEM. Statistical tests: one-way ANOVA and Tukey post-hoc; ns: *p* > 0.05, \**p* < 0.05, \*\*\**p* < 0.001.

## DISCUSSION

Regulation of energy intake is a complex process involving food search, an animal’s internal state, and the sensory qualities of food. Fructose, either consumed directly or rapidly metabolized from precursors, promotes feeding through activation of a brain fructose sensor called Gr43a (Miyamoto et al., 2012). Here we describe how a neuronal network composed of neurons in the Fan-shaped body and asymmetric body contributes to energy homeostasis by detecting satiety-dependent changes in hemolymph glucose and modulating fructose drive (Figure 8C).

### The FB is a very organized yet incompletely understood structure

The central complex, which is comprised of the FB, the protocerebral bridge (PB), the ellipsoid body, and the noduli, is regarded as a center for sensory motor integration that functions in goal directed behavior (Fisher et al., 2019; Honkanen et al., 2019; Hulse et al., 2020; Pfeiffer and Homberg, 2014; Sun et al., 2020; Weir and Dickinson, 2015; Wolff et al., 2015). The FB is organized in nine horizontal layers and nine vertical columns. FB large field neurons of layers 1 to 3, and inputs to these layers from the PB, encode flight direction and general sensory orientation (Currier et al., 2020; Weir and Dickinson, 2015). FB layers 6 and 7 are well known to regulate sleep and arousal (Berry et al., 2015; Donlea et al., 2011; Ueno et al., 2012), locomotor control (Strauss, 2002), courtship (Sakai and Kitamoto, 2006) and visual memory (Li et al., 2009; Liu et al., 2006; Wang et al., 2008). Layer 6 also plays a role in avoiding conditioned odors, while layers 1,2, 4 and 5 respond to electric stimuli and are required for innate odor avoidance (Hu et al., 2018). However, the function of the most dorsal FB layers (8 and 9), mostly innervated local tangential neurons and AB-FBl8 (or vΔA_a), remained poorly understood (Hulse et al., 2020; Jenett et al., 2012; Wolff and Rubin, 2018; Wolff et al., 2015). Our results demonstrate an involvement of these layers in feeding regulation.

### SLP-AB and AB-FBl8 provide insight on asymmetric body function

We find that AB-FBl8 oscillations require glutamatergic input from SLP-AB projections to the asymmetric body. Described for the first time in 2004 (Pascual et al., 2004), very little is known about the asymmetric body. 92.4% of flies display asymmetry in this structure, with the body present only in the right hemisphere, while 7.6% also have a body on the left side (Pascual et al., 2004). We noted that oscillations in the AB-FBl8 display a tendency to be faster on the right side, with clearly asynchronous activity that may reflect their asymmetric input from SLP-AB. Interestingly, the small proportion of flies displaying symmetry have defects in long-term memory, a process that is known to require energy (Burke and Waddell, 2011; Mery and Kawecki, 2005; Musso et al., 2015; Pascual et al., 2004; Plaçais and Preat, 2013; Plaçais et al., 2017). We speculate that these symmetric flies may have a dysfunctional SLP-AB to AB-FBl8 connection, resulting in impaired Tk release. This could impact LTM either directly or through changes in feeding (Nässel et al., 2019). A role for TK in memory has been demonstrated in honeybees (Boerjan et al., 2010; Brockmann et al., 2009; Takeuchi et al., 2004) and mammals (Lénárd et al., 2018), and TkR86C appears to be expressed in serotonergic paired neurons (SPN) known to interact with MB-MP1 neurons required for long-term memory formation (Poels et al., 2009; Scheunemann et al., 2018). Tk also acts through TkR99D to modulate activity in neurons producing i-like peptides (Birse et al., 2011; Qi et al., 2020), which impact LTM formation (Chambers et al., 2015; Eschment et al., 2020).

Surprisingly, modulation of AB-FBl8 oscillations by SLP-AB requires glutamatergic signaling through GluClα. This suggests that either inhibition from SLP-AB is required for AB-FBl8 oscillatory activity or AB-FBl8 have an unusual chloride reversal potential. Interestingly, GluClα has been previously implicated in ON/OFF responses of the visual system, demonstrating a role in regulating cell excitability (Molina-Obando et al., 2019). Further study will be required to fully understand this connection, along with the source of input to SLP-AB in the protocerebrum.

### A dual role for 2 carbohydrates

Since glucose is the primary circulating energy source, one might intuitively expect that enhancing feeding in response to post-ingestive glucose detection would be the most efficient means of energy uptake. However, using elevation of hemolymph glucose as a signal to continue feeding is problematic because glucose levels are tightly regulated, and elevated glucose serves as a signal of satiety. On the other hand, circulating fructose can vary widely in response to ingestion and can therefore be a more reliable indicator of recent sugar intake (Miyamoto et al., 2012). Moreover, fructose typically co-exists with other nutritive sugars in common food sources and can therefore serve as an effective proxy for carbohydrate ingestion. Thus, the separation of glucose as a satiety indicator and fructose as marker of sugar consumption removes the potential ambiguity of each as a signal. To enable sustained feeding on a rich sugar source, such a mechanism would require that ingestion of sugars and the subsequent activation of Gr43a brain neurons rapidly promotes feeding behavior. Our results support this idea in two main ways. First, silencing of AB-FBl8 neurons by genetic manipulation or prolonged starvation produces Gr43a-dependent fructose preference within the first 10 minutes of a FlyPAD assay (Figure S1C). Second, closed-loop optogenetic activation of Gr43a brain neurons was sufficient to produce a strong positive preference within 60 minutes in the STROBE (Figure 8B).

Interestingly, glucose and fructose also have differential effects in mammals, where glucose ingestion promotes satiety by repressing the secretion of the hunger hormone ghrelin and stimulating the secretion of satiety hormones such as leptin or insulin (Merino et al., 2019). Fructose, however, induces a reduction in ghrelin and can promote over-feeding (Merino et al., 2019). Similarly, hypothalamic AMPK is described as functioning as an energy sensor and its inhibition by glucose administration or leptin and insulin promotes anorexigenic behavior. Conversely, fructose administration activates AMPK and promotes orexigenic behavior (Burmeister et al., 2013; Cawley, 2012; Cha et al., 2008; Minokoshi et al., 2004; Wang et al., 2004a). The first description of fly Gr43a neurons noted their orexinegenic activity and suggested a potential functional homology with the hypothalamus (Miyamoto et al., 2012). In the present study, we uncovered a multi-layered neural system centered on a brain energy sensor (AB-FBl8), whose activation by glucose leads to anorexigenic behavior through inhibition of the brain fructose sensor Gr43a. Thus, our results are consistent with at least partial functional homology between the mammalian hypothalamus and brain Gr43a neurons of the fly.

## Supporting information

raw data for figure 1

raw data for figure 2

raw data for figure 3

raw data for figure 4

raw data for figure 5

raw data for figure 6

raw data for figure 7

raw data for figure 8

raw data for figure S1

raw data for figure S2

raw data for figure S3

raw data for figure S4

raw data for figure S5

raw data for figure S6

raw data for figure S7

raw data for figure S8

## SUPPLEMENTAL INFORMATION

Supplemental information includes eight figures that can be found in an associated file, and two supplemental movies.

## ACKNOWLEDGEMENTS

We thank members of the lab for their comments on the manuscript. We thank Dr. Pierre-Yves Plaçais for providing the MATLAB code for oscillation analysis and for his help in processing. We also thank Hubert Amrein, the Bloomington stock center, and the Vienna *Drosophila* Resource Center for fly stocks. This work was funded by the Natural Sciences and Engineering Research Council (NSERC) of Canada (RGPIN-2016-03857 and RGPAS-49246-16) and a Michael Smith Foundation for Health Research Scholar Award to M.D.G.

## AUTHOR CONTRIBUTIONS

Conceptualization, P-Y.M; Methodology, P-Y.M and M.D.G; Formal analysis, P-Y.M, P.J, and M.D.G; Investigation, P-Y.M, and P.J; Writing, P-Y.M and M.D.G; Visualization, P-Y.M; Supervision, M.D.G; Funding acquisition, M.D.G.

## DECLARATION OF INTERESTS

Authors declare no conflicts of interests.

## KEY RESSOURCES TABLE

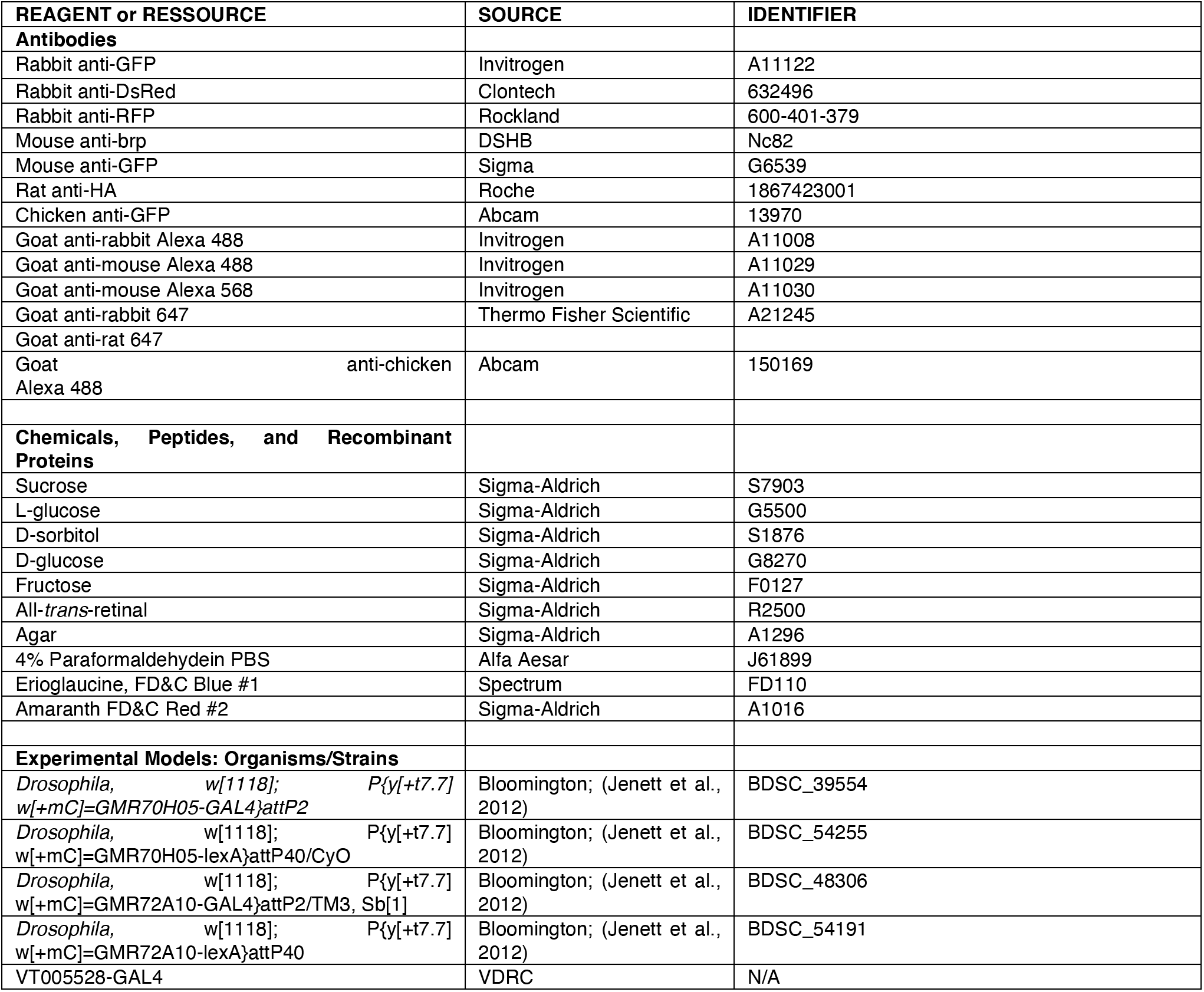
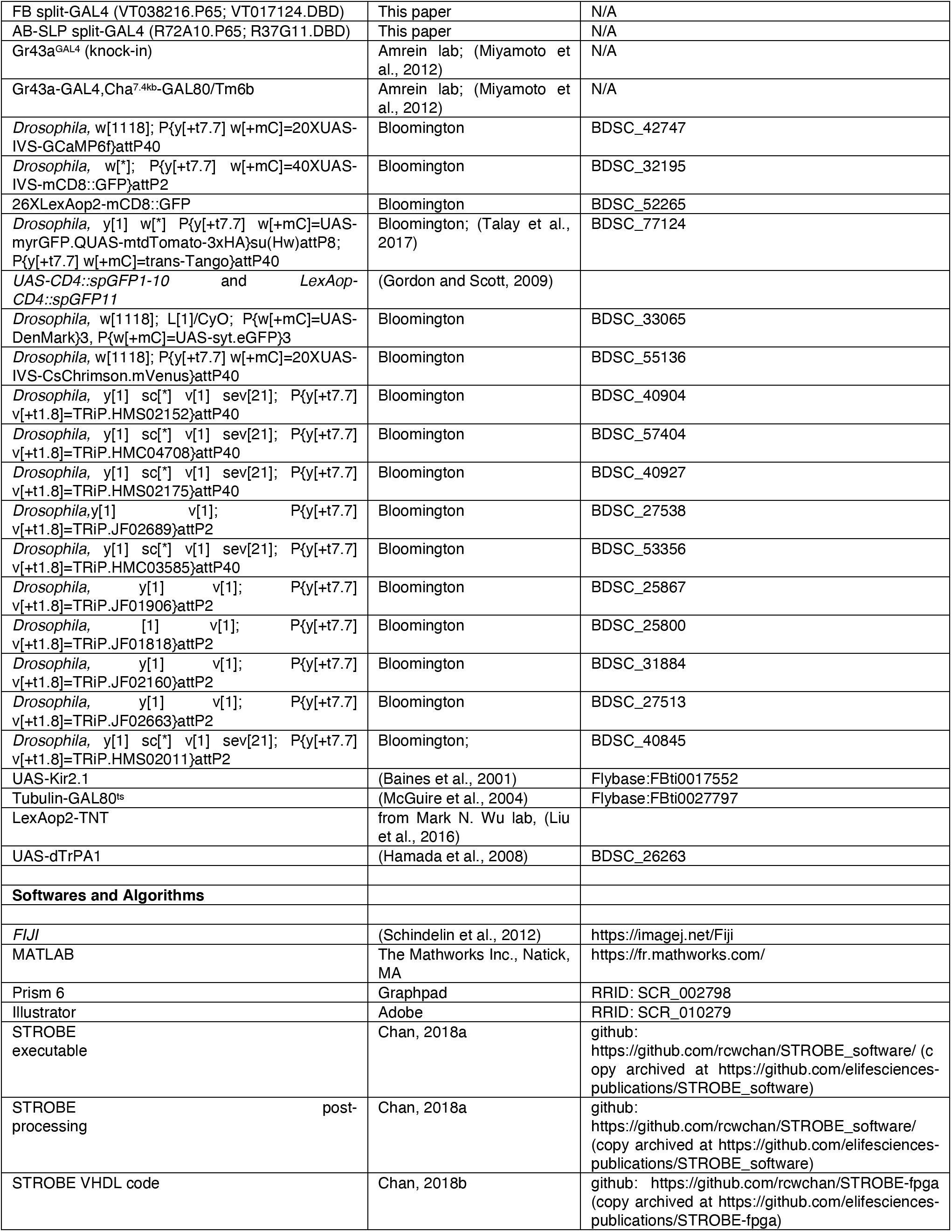

## RESOURCE AVAILABILITY

### Lead Contact

Further information and requests for resources and reagents should be directed to and will be fulfilled by the Lead Contact, Dr Micheal D. Gordon.

There are no restrictions on reagent sharing to disclose

### Materials Availability

Flies generated in this study are available upon request.

### Data and Code Availability

All STROBE software is available for download from Github: FPGA code: https://github.com/rcwchan/STROBE-fpga (copy archived at https://github.com/elifesciences-publications/STROBE-fpga).

All other code: https://github.com/rcwchan/STROBE_software/ (copy archived at https://github.com/elifesciences-publications/STROBE_software).

## EXPERIMENTAL MODEL AND SUBJECT DETAILS

### Drosophila melanogaster

Fly stocks were raised on standard food at 25°C and 70% relative humidity under a 12:12 hr light:dark cycle. For neuronal silencing experiments we used *UAS-Kir2.1* (Baines et al., 2001), *tub-Gal80*^ts^ (McGuire et al., 2004), and *LexAop-tnt* (Liu et al., 2016). For neuronal activation experiments we used *UAS-dTrpA1* (Hamada et al., 2008) and *20XUAS-IVS-CsChrimson.mVenus* (Bloomington #55135). Specific FB expression was driven using *R70H05-GAL4* ((Jenett et al., 2012); Bloomington #39554), *R70H05-LexA* ((Jenett et al., 2012); Bloomington #54255), *VT005528-GAL4* (VDRC) and a newly built split-Gal4 line (*VT038216.P65; VT017124.DBD*). Specific SLP-AB expression was driven using *R72A10-GAL4* ((Jenett et al., 2012); Bloomington #48306), *R72A10-LexA* ((Jenett et al., 2012); Bloomington #54191) and a newly built split-GAL4 line (*R72A10.P65; R37G11.DBD*). Specific Gr43a expression was driven using *Gr43a^GAL4^* (knock-in) and *Gr43aGal4,Cha^7.4kb^-GAL80* ((Miyamoto et al., 2012); gift from H. Amrein). For staining experiments, we used *40XUAS-IVS-mCD8∷GFP* (Bloomington #32195), *26XLexAop2-mCD8∷GFP* (Bloomington #77124),*Trans-Tango* ((Talay et al., 2017); Bloomington #77124), *20XUAS-IVS-CsChrimson* (Bloomington #55136), and *UAS-DenMark,UAS-Syt* (Nicola et al., - Bloomington #33065). For GRASP experiment, we used *UAS-CD4∷spGFP1-10* and *LexAop-CD4∷spGFP11* (Gordon and Scott, 2009). For imaging experiments we used the *20XUAS-IVS-GCaMP6f* (Bloomington #42747). For RNAi experiments, we used RNAi against Glut1 (Bloomington #40904), HexC (Bloomington #57404), Vglut (Bloomington #40927), Vglut (Bloomington #27538), Vglut (Bloomington #40845) GluCla (Bloomington #53356), sNPF (Bloomington v25867), Tk (Bloomington #25800), TkR 99D (Bloomington #27513), and TkR86C (Bloomington #31884).

## METHOD DETAILS

### Fly preparation and behavior experiments

All experiments were performed with mated female flies to reduce variability, given that sex differences were not a subject of investigation. After eclosion, flies were kept for 2 to three days in fresh vials containing standard medium. For thermo-sensitive silencing experiments (Kir^ts^), flies were then transferred into vials for 2 days at 29 °C. Flies were subjected to a varying fasting period (0-30 hrs) where they were transferred to vials containing 1 ml of 1% agar at 29 °C. For silencing (*UAS-kir2.1*; *LexAop-tnt*) and RNAi experiments, flies were transferred into vials containing 1 ml of 1% agar at 29 °C for 15-18 hrs. For activation experiments (dTrpA1), flies were transferred into vials containing 1 ml of 1% agar at 22 °C for 43-45 hrs. For STROBE experiments, flies were kept for several days in fresh vials containing standard medium, and were then transferred at 25°C into vials covered with aluminum foil containing 1 ml standard medium (control flies) or 1 ml standard medium containing with 1 mM of all-*trans*-retinal (retinal flies) for 2 days. Flies were then subjected to a 24-hr fasting period where they were transferred to covered vials containing 1 ml of 1% agar (control flies) or 1 ml of 1% agar mixed with 1 mM of all-*trans*-retinal (retinal flies). Sucrose, L-glucose, D-glucose, D-sorbitol, D-fructose and agar were obtained from Sigma-Aldrich.

### FlyPAD experiments

All flies were 5-9 days old at the time of the assay, and experiments were performed between 10:00 am and 5:00 pm. For single tastant experiments, one channel of the arena was loaded with 3.5 μl of 1% agar Mixed with a tastant. To exclude interactions due to drinking behavior, the other side was loaded with 3.5 μl of 1% agar. The tastant used were sucrose (5, 50 and 1000 mM); L-glucose (5, 50 and 1000 mM), L-glucose 50 mM combined with D-sorbitol (0, 5, 50, 500 and 1000 mM); D-glucose (5, 50 and 1000 mM); fructose (5, 50 and 1000 mM); and D-sorbitol (50 mM). For dual tastant experiments, one channel was loaded with fructose while the other one with D-glucose, always in an equimolar manner (5, 50 and 1000 mM). Acquisition on the FlyPAD software was started and then single flies were transferred into each arena by mouth aspiration. Experiments were run for 60 min, and the preference index for each fly was calculated as: (interactions with Food 1 – interactions with Food 2)/(interactions with Food 1 + interactions with from Food 2). Tastants were all obtained from Sigma-Aldrich.

### STROBE experiments

Experiments were performed as previously described (Jaeger et al., 2018; Musso et al., 2019). All flies were 5-9 days old at the time of the assay, and experiments were performed between 10:00 am and 5:00 pm. Both channels of the arena were loaded with 3.5 μl of 1% agar. Acquisition on the STROBE software was started and then single flies were transferred into each arena by mouth aspiration. Experiments were run for 60 min, and the preference index for each fly was calculated as: (interactions with Food 1 - interactions with Food 2)/(interactions with Food 1 + interactions with from Food 2). The red LED is always associated to the left side (Food 1), with a light intensity of 11.2 mW/cm^2^. Agar and all-*trans*-retinal were obtained from Sigma-Aldrich.

### 2-Choice

Binary choice preference tests were similar to those previously described (Jaeger et al., 2018; LeDue et al., 2015). Female flies aged 2–5 days were sorted into groups of 10 and were transferred starved as explained above. For the assays, flies were then transferred into testing vials containing six 10 μL dots of agar that alternated in color. The food choices were: 1% agar with 50 mM fructose (Food 1), and 1% agar with 50 D-glucose (Food 2). Each choice contained either 0.125 mg/mL blue (Erioglaucine, FD and C Blue#1) or 0.5 mg/mL red (Amaranth, FD and C Red#2) dye, and half the replicates for each experiment were done with the dyes swapped to control for any dye preference. Flies were allowed to feed for 2 hr in the dark at 29°C and then frozen and scored for abdomen color. Preference index (PI) was calculated as ((# of flies labeled with Food 1 color) − (# of flies labeled with Food 2 color))/(total number of flies that fed).

### PER

For tarsal PER, flies were mounted on glass slides using nail polish. For labellar PER, flies were placed inside a pipette tip cut to size so that only the head was exposed. Flies were then sealed into the tube with tape, and then adhered to a glass slide with double-sided tape. Flies were allowed 1–2 h to recover before testing began. Flies were stimulated with water on their front tarsi or labella for tarsal and labellar PER, respectively, and allowed to drink until satiated. Each fly was then stimulated with increasing concentration of a either D-glucose or fructose on either the tarsi or labella, and responses to each tastant were recorded. Flies were provided with water between each tastant. All stimuli were delivered with a 1 ml syringe attached to a 20 μl pipette tip.

### *In-Vivo* Calcium imaging

Female flies aged 5-9 days were briefly anesthetized. With a custom chamber, each fly was mounted by insertion of the cervix into individual collars. For further immobilization of the head, nail polish was applied in a thin layer to seal the head to the chamber. The antennae and the associated cuticle covering the SEZ were removed until the ocelli, and adult hemolymph-like (AHL) buffer with ribose (Liu et al., 2012; Wang et al., 2003) was immediately injected into the preparation to cover the exposed brain. Flies were left to recover from anesthesia for an hour before imaging. At the beginning and end of the experiment, spontaneous or brush tickling–evoked leg or abdomen movement was checked to ensure that the fly was still alive.

GCaMP6f fluorescence was imaged with a Leica SP5 II laser scanning confocal microscope equipped with a tandem scanner and HyD detector. The relevant area of the FB was visualized using the 25× water objective. Images were acquired at a speed of 8,000 lines per second with a line average of 1, resulting in a collection time of 0.051 ms per frame at a resolution of 256× 126 pixels for a total of 7 minutes. The pinhole was opened to 200 μm.

Image analysis was performed following a previously described protocol (Musso et al., 2015; Plaçais and Preat, 2013; Plaçais et al., 2012). It was performed offline with a custom-written Matlab program. Light intensity was averaged over a region of interest delimited by hand and surrounding the projections of AB-FBl8 neurons on the FB layers 8 and 9. Three areas of interest (ROI) were analyzed: the tips and the central part. From a given region of interest, the resulting time trace was normalized to a percent change of fluorescence (100 (*F* − *F*_0_) / *F*_0_), using a baseline value of the fluorescence *F*_0_ that was estimated as the mean fluorescence over the whole acquisition. To remove long-term drift, a baseline resulting from the moving average over a 100-s time window was then subtracted from the signal. Thus, in subsequent frequency analyses, all frequency axes are presented starting at 0.01 Hz. Given that signals are noisy, their amplitudes were estimated as the difference between the means of the 30% upper and lower quantiles of data points. For each signal, the power spectrum was computed and smoothed over a frequency window of 0.02 Hz. Rhythmic spontaneous activity in the time domain resulted in a peak in the power spectrum that had a finite width, as oscillations are intrinsically noisy. A fit of a Lorentzian curve to the power spectrum was performed to yield an estimate of the central frequency of the peak, *f*_0_, and the width of the peak at half its maximal value, Δ*f*. *f*_0_ defined the characteristic frequency of the oscillation and frequency fluctuations around *f*_0_, and hence the regularity of the oscillation, could be quantified by the quality factor *Q* = *f*_0_/Δ*f* (Plaçais et al., 2009). A quality factor greater than 0.5 indicates that the zero frequency is excluded from the peak: this value was thus taken as a threshold to define a signal as rhythmically oscillating. When the fitting procedure converged to a value below 0.5, it was thus irrelevant to define oscillating parameters, and *f*_0_ and *Q* were both assigned zero values.

To plot average amplitude histograms, we calculated a mean amplitude value for the different ROI selected, and then averaged the mean values across all flies from the same condition.

Average power spectra across all animals from the same condition were obtained and were additionally smoothed over a 0.03-Hz frequency window. Peaked average spectra (Figure 1D, Figure S1A-B, Figure S2A-C, Figure 4G, Figure S4A-B, Figure 5D, Figure S6G-H) were characterized by their mean frequency *f*_0_ and a quality factor *Q* calculated from *f*_0_ and the width at half-height.

### Immunohistochemistry

For GFP, Brain immunofluorescence was carried out as described previously (Chu et al., 2014). Primary antibodies used were Rabbit anti-GFP (1:1000, Invitrogen) and mouse anti-nc82 (1:50, DSHB). Secondary antibodies used were goat anti-Rabbit Alexa 488 (1:200, Invitrogen) and goat anti-mouse Alexa 568 (1:200, Invitrogen). For DenMark,Syt immunofluorescence, primary antibodies used were chicken anti-GFP (1:1000, Abcam), and rabbit anti-RFP (1:200, Rockland). Secondary antibodies used were goat anti-chicken Alexa 488 (1:200, Abcam) and goat anti-rabbit Alexa 647 (1:200, Thermo Fisher Scientific, Waltham, MA, #A21245). For GRASP immunofluorescence, primary antibodies used were Mouse anti-GFP (1:100, Sigma, catalog #G6539) and rabbit anti-DsRed (1:2,000, Clontech, #632496). Secondary antibodies used were goat anti-Mouse Alexa 488 (1:200, Invitrogen) and goat anti-Rat Alexa 568 (1:200, Invitrogen). For Trans-Tango immunofluorescence, primary antobodies were Rabbit anti-GFP (1:1000, Invitrogen), mouse anti-nc82 (1:50, DSHB), and rat anti-HA (1:100, Roche). Secondary antibody used were goat anti-Rabbit Alexa 488 (1:200, Invitrogen), goat anti-mouse Alexa 568 (1:200, Invitrogen), and goat anti-rat Alexa 647 (1:200, Invitrogen). All images were acquired using a Leica SP5 II Confocal microscope with a 25x water immersion objective. All images were taken sequentially with a z-stack step size at 1 μm, a line average of 2, line-scanning speed of 200 Hz, and a resolution of 1024 × 1024 pixels. Images were processed in ImageJ (Schneider et al 2012).

## QUANTIFICATION AND STATISTICAL ANALYSIS

Statistical tests were performed using GraphPad Prism six software. Descriptions and results of each test are provided in the figure legends. Sample sizes are indicated in the figure legends. Sample sizes were determined prior to experimentation based on the variance and effect sizes seen in prior experiments of similar types. Whenever possible, all experimental conditions were run in parallel and therefore have the same or similar sample sizes.

All replicates were biological replicates using different flies. Data for all quantitative experiments were collected on at least three different days, and behavioral experiments were performed with flies from at least two independent crosses. Specific definitions of replicates are as follows. For calcium imaging, each data point represents the activity of a single fly to the indicated condition. For binary choice behavioral tests, each data point represents the calculated preference for a group of 10 flies. For PER, each replicate is composed of 10 independent flies tested in parallel. For flyPAD and STROBE experiments, each data point is the calculated preference of an individual fly over the course of the experiment.

There were two conditions where data were excluded that were determined prior to experimentation and applied uniformly throughout. First, in calcium imaging experiments, all the data from a fly were removed if either: a) there was too much movement during the recording to reliably quantify the response; or b) flie were dead at the end of the recording. Second, for flyPAD and STROBE experiments, the data from individual flies were removed if the fly did not pass a set minimum threshold of sips (10), or the data showed hallmarks of a technical malfunction (rare).

All the quantitative data used for statistical tests can be found as supplements for each figure.

## Notes

### Competing Interest Statement

The authors have declared no competing interest.

