## Supplemental figures for "A neural circuit linking two sugar sensors regulates satiety-dependent fructose drive in *Drosophila*"

### SUPPLEMENTAL INFORMATION

**Supplemental Movie 1. Strong calcium activity of fed *R70H05-GAL4 > UAS-GCaMP6f* flies, related to Figure 1**

**Supplemental Movie 2. Weak calcium activity of 30 hrs starved *R70H05-GAL4 > UAS-GCaMP6f* flies, related to Figure 1**

### SUPPLEMENTAL FIGURES

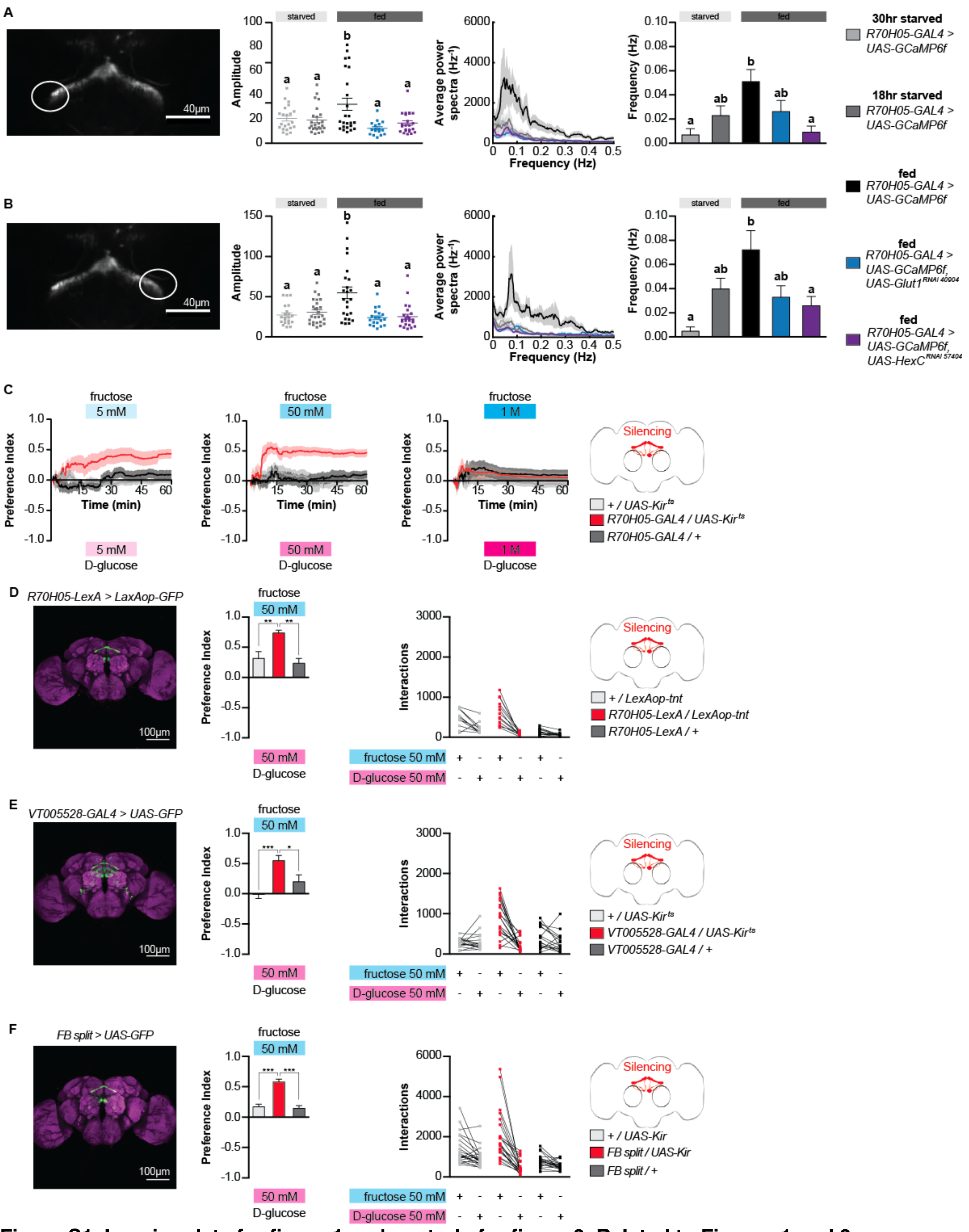


**Figure S1. Imaging data for figure 1 and controls for figure 3, Related to Figures 1 and 3**

(A) Imaging data from left tip of AB-FBl8 neurons (*n* = 17-25). (B) Imaging data from right tip of AB-FBl8 neurons (*n* = 19-25). (C) Preference index for *R70H05-GAL4 > UAS-Kir2.1-GAL80$^{ts}$* flies over the course of a 1 hr experiment from figure 3B. (D) Immunofluorescent detection of *LexAop-GFP* driven by *R70H05-LexA* (left); preference Index (middle) and Interactions of *R70H05-LexA > LexAop-tnt* flies between 50 mM fructose and D-glucose (right; *n* = 10-14). (E) Immunofluorescent detection of UAS-GFP driven by VT005538-GAL4 (left); preference index (middle) and Interactions of *VT005528-GAL4 > UAS-Kir2.1-GAL80$^{ts}$* flies between 50 mM fructose and D-glucose (right; *n* = 16-20). (F) Immunofluorescent detection of *UAS-GFP* driven by *VT038216.P65;VT017124.DBD* (left); preference Index (middle) and Interactions of *VT038216.P65;VT017124.DBD > UAS-Kir2.1* flies between 50 mM fructose and D-glucose (right; *n* = 21-23). Values represent mean ± SEM. Statistical tests: one-way ANOVA and Tukey post-hoc; ns: $p > 0.05$, * $p < 0.05$, ** $p < 0.01$, ***$p<0.001$.

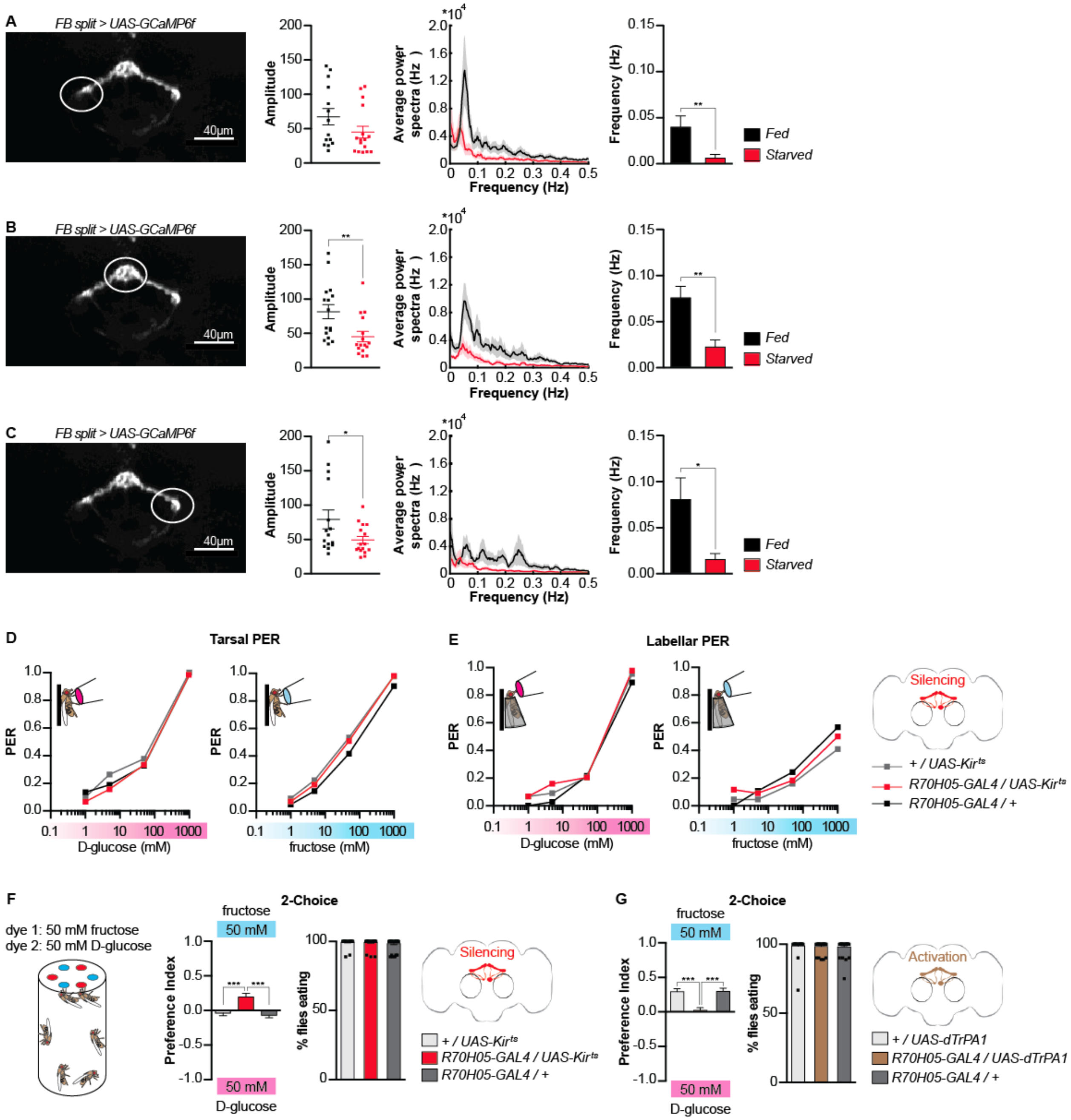


**Figure S2. Silencing AB-FBl8 modifies fructose ingestion, Related to Figure 3**

(A-C) Flies expressing *UAS-GCaMP6f* under control of the *FB-split* driver display strong oscillatory activity when fed, which decreases following starvation (40 hours of starvation; A: $n$ = 14-16; B: $n$ = 16; C: $n$ =15-16). (D) Tarsal PER of AB-FBl8-silenced flies to D-glucose (left; $n$ = 39-58) and fructose (right; $n$ = 43-58). (E) Labellar PER of AB-FBl8- silenced flies to D-glucose (left; $n$ = 37-44) and fructose (right; $n$ = 37-44). (F) Effect of silencing AB-FBl8 neurons on the 2-choice ingestion preference between 50 mM fructose and D-glucose, following 16 hours of starvation ($n$ = 39-40. (G) Effect of activating AB-FBl8 neurons on the 2-choice ingestion preference between 50 mM of fructose and D-glucose, following 43 hours of starvation ($n$ = 34-35). For PER, values are the fraction of flies that responded; For 2-choice values represent mean ± SEM. Statistical tests: one-way ANOVA and Tukey post-hoc for behavior and t-test for imaging; ns: $p > 0.05$, * $p < 0.05$, ** $p < 0.01$, ***$p<0.001$.

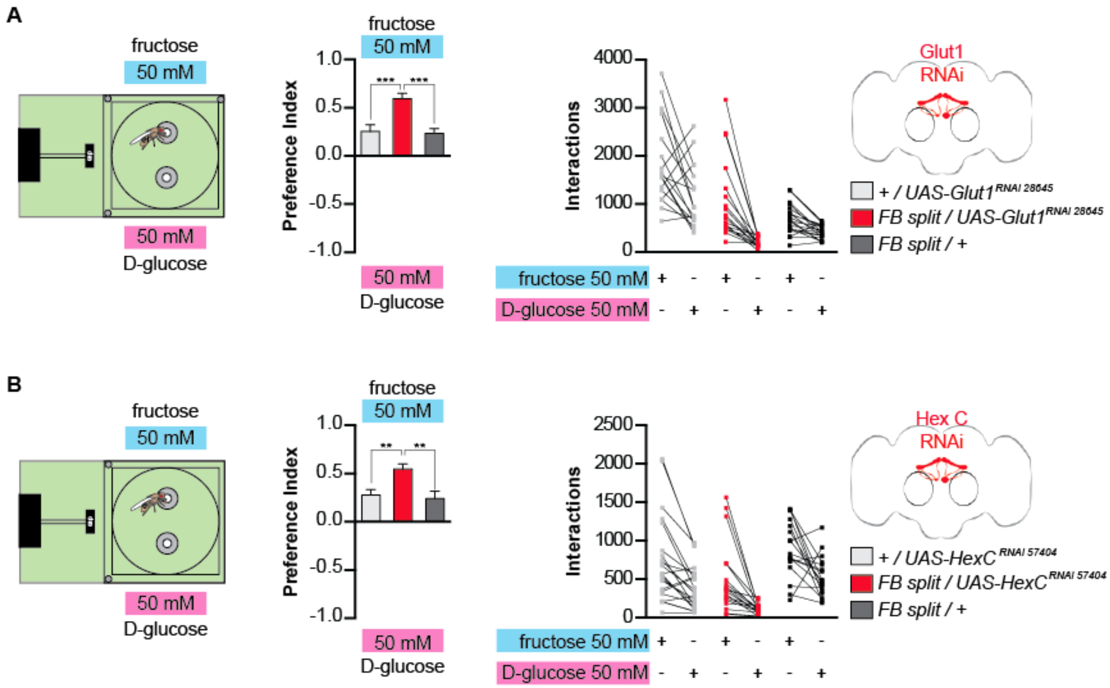


**Figure S3. Modulation of fructose feeding preference requires the expression of Glucose transporter 1 and Hexokinase C in the AB-FBl8 neurons, Related to Figure 3**

(A) Effect of knocking down Glut1 in AB-FBl8 neurons on the preference between 50 mM fructose and D-glucose (B: $n$ = 18-20). (B) Effect of knocking down HexC in AB-FBl8 neurons on the preference between 50 mM fructose and D-glucose (D: $n$ = 17-22). Values represent mean ± SEM. Statistical tests: one-way ANOVA and Tukey post-hoc; ns: $p > 0.05$, ** $p < 0.01$, ***$p<0.001$.

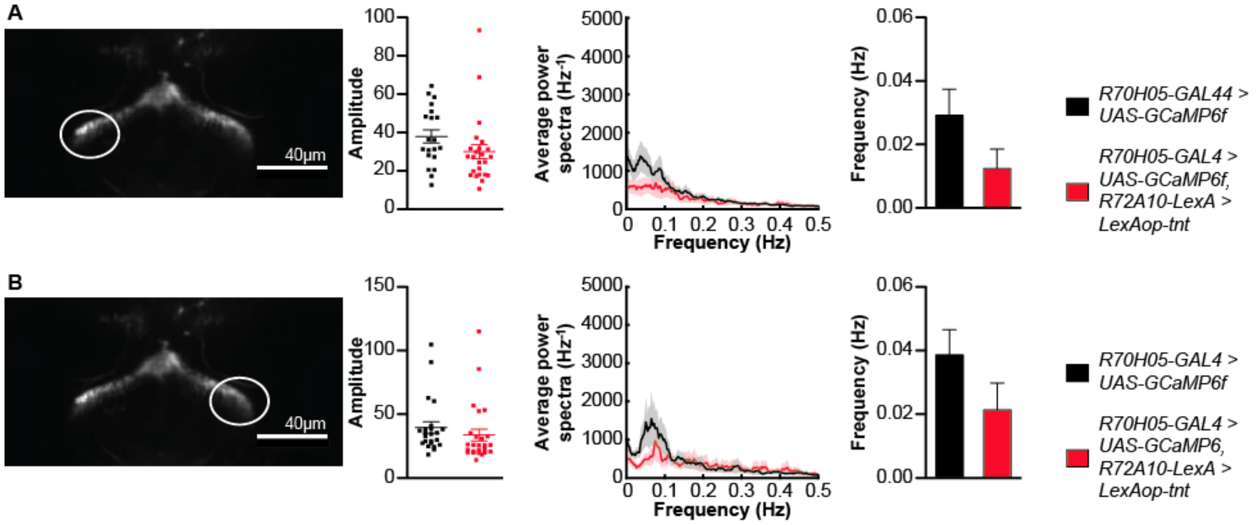


**Figure S4. Imaging data from SLP-AB silenced flies, Related to Figure 4**
(A-B) Silencing SLP-AB neurons does not affect oscillations in the tips of AB-FBl8 neurons (A: $n$ = 20-24; B: $n$ = 23-25). Values represent mean ± SEM. Statistical tests: *t*-test.

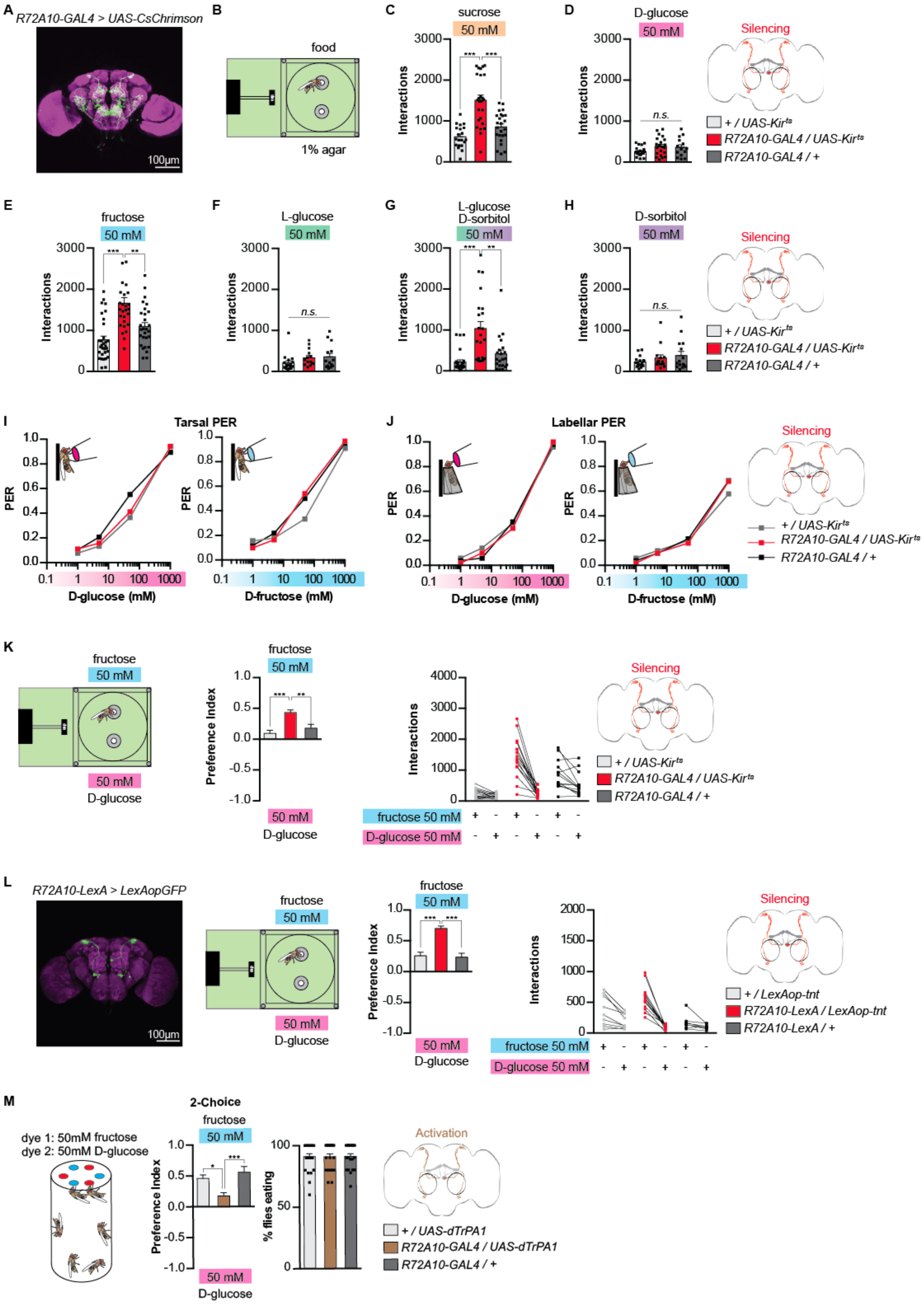
A
R72A10-GAL4 > UAS-CsChrimson
100μm
B
food
1% agar
C
sucrose
50 mM
Interactions
D
D-glucose
50 mM
n.s.
Silencing
+ / UAS-Kir^ts
R72A10-GAL4 / UAS-Kir^ts
R72A10-GAL4 / +
E
fructose
F
L-glucose
G
L-glucose
D-sorbitol
H
D-sorbitol
I
Tarsal PER
J
Labellar PER
PER
D-glucose (mM)
D-fructose (mM)
K
Preference Index
fructose 50 mM
D-glucose 50 mM
L
R72A10-LexA > LexAopGFP
+ / LexAop-tnt
R72A10-LexA / LexAop-tnt
R72A10-LexA / +
M
2-Choice
dye 1: 50mM fructose
dye 2: 50mM D-glucose
% flies eating
Activation
+ / UAS-dTrPA1
R72A10-GAL4 / UAS-dTrPA1
R72A10-GAL4 / +

**Figure S5. Silencing SLP-AB neurons increases fructose feeding preference, Related to Figure 5**
(A) Immunofluorescent detection of *UAS-Chrimson* driven by *R72A10-GAL4*. (B) Experimental setup: one channel is filled with food and the other one is filled with 1% agar. (C-H) Effect of silencing SLP-AB neurons on flies' interactions with 50 mM of sucrose (C; $n$ = 20-25)), 50 mM of D-glucose (D; $n$ = 14-21), 50 mM of fructose (E; $n$ = 28-31)), 50 mM of L-glucose (F; $n$ = 14-17), 50 mM of L-glucose and D-sorbitol (G; $n$ = 21-27), and 50 mM of D-sorbitol (H; $n$ = 8-11). (I) Tarsal PER of SLP-AB-silenced flies to D-glucose (left; $n$ = 52-64) and fructose (right; $n$ = 45-63). (J) Labellar PER of SLP-AB-silenced flies to D-glucose (left; $n$ = 45-51) and fructose (right; $n$ = 50). (K) Effect of silencing SLP-AB neurons on flies' preference between fructose and D-glucose, and their corresponding interactions ($n$ = 14-17). (L) Immunofluorescent detection of *LexAop-GFP* driven by *R72A10-LexA*; and effect of silencing these neurons on flies' preference between fructose and D-glucose, and their corresponding interactions ($n$ = 8 13). (M) Effect of activating SLP-AB neurons on the 2-choice ingestion preference between 50 mM fructose and D-glucose, following 43 hours of starvation ($n$ = 20-24). For PER, values are the fraction of flies that responded; For 2-choice and flyPAD, values represent mean ± SEM. Statistical tests: one-way ANOVA and Tukey post-hoc for behavior and t-test for imaging; ns: $p > 0.05$, * $p < 0.05$, ** $p < 0.01$, ***$p<0.001$.

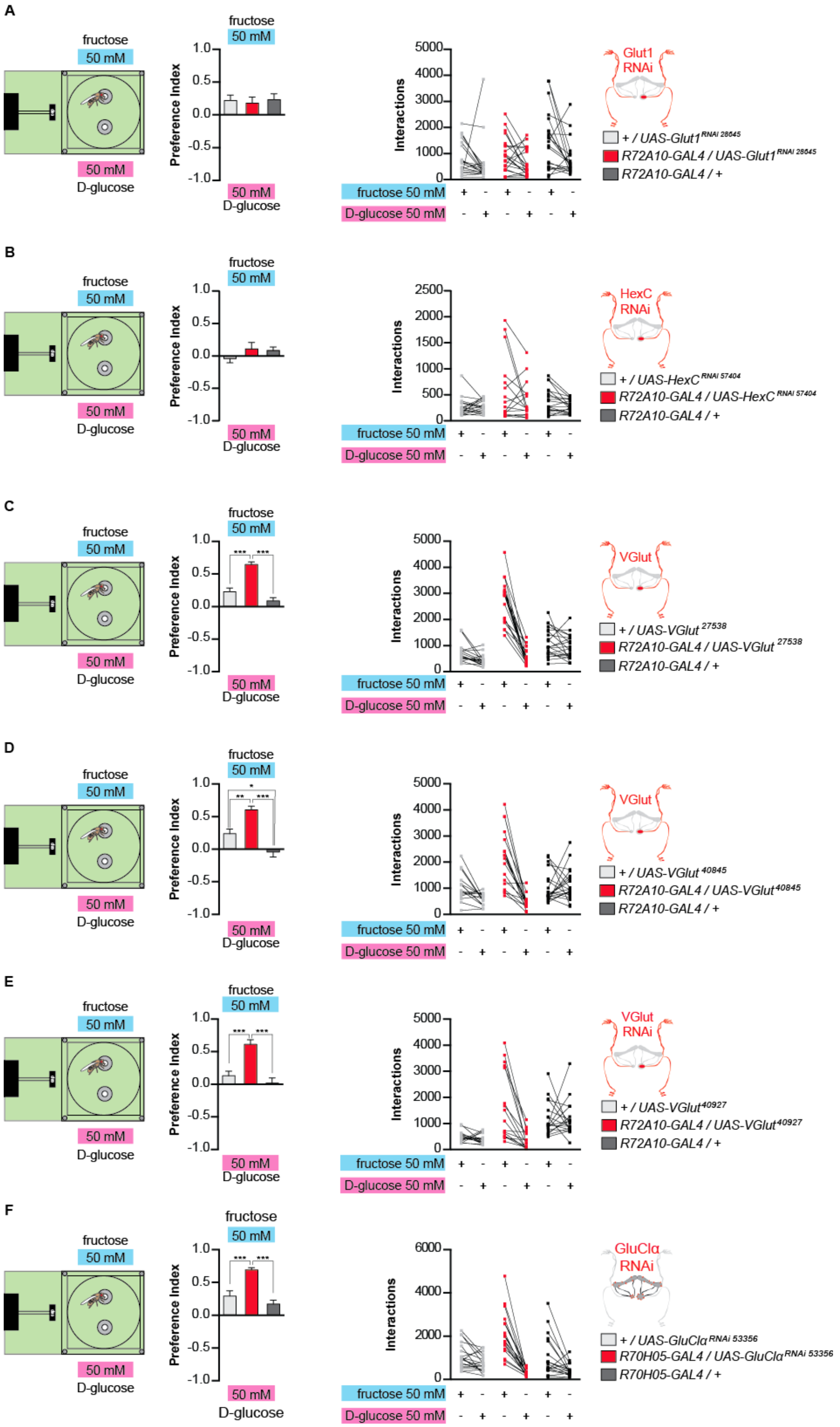
A
fructose
50 mM
50 mM
D-glucose
Preference Index
Interactions
fructose 50 mM
D-glucose 50 mM
Glut1
RNAi
+ / UAS-Glut1 RNAi 28645
R72A10-GAL4 / UAS-Glut1 RNAi 28645
R72A10-GAL4 / +
B
HexC
RNAi
+ / UAS-HexC RNAi 57404
R72A10-GAL4 / UAS-HexC RNAi 57404
R72A10-GAL4 / +
C
VGlut
+ / UAS-VGlut 27538
R72A10-GAL4 / UAS-VGlut 27538
R72A10-GAL4 / +
D
VGlut
+ / UAS-VGlut 40845
R72A10-GAL4 / UAS-VGlut 40845
R72A10-GAL4 / +
E
VGlut
RNAi
+ / UAS-VGlut 40927
R72A10-GAL4 / UAS-VGlut 40927
R72A10-GAL4 / +
F
GluClα
RNAi
+ / UAS-GluClα RNAi 53356
R70H05-GAL4 / UAS-GluClα RNAi 53356
R70H05-GAL4 / +

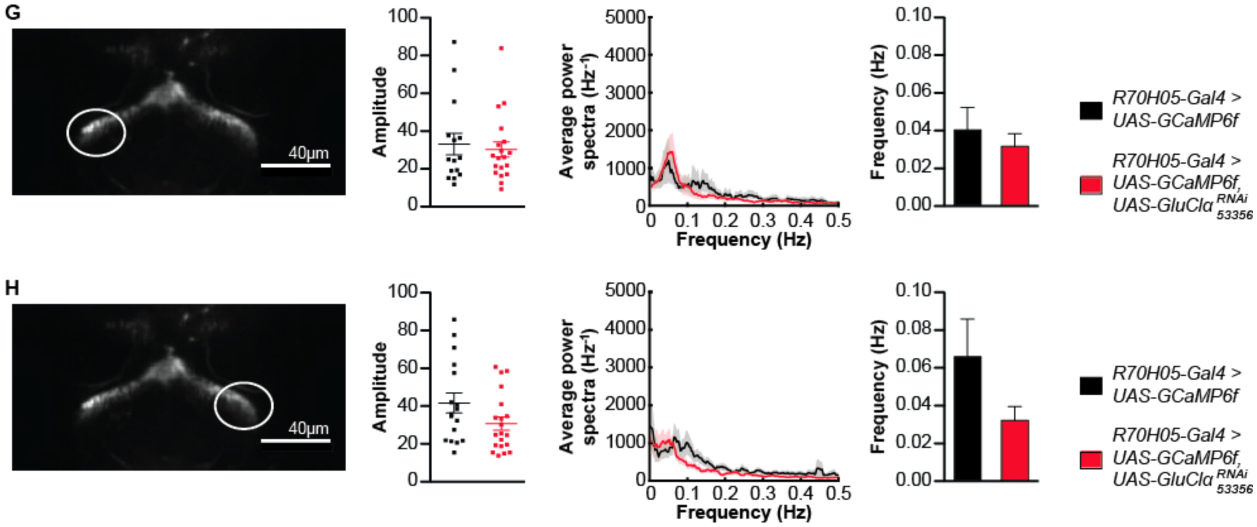


**Figure S6. Glutamatergic SLP-AB neurons are positively connected to AB-FBl8 neurons, related to Figure 5**

(A) Effect of knocking down Glut1 in SLP-AB neurons on flies' preference between fructose and D-glucose, and their corresponding interactions ($n$ = 20-21). (B) Effect of knocking down HexC in SLP-AB neurons on flies' preference between fructose and D-glucose, and their corresponding interactions ($n$ = 16-20). (C-E) Effect of knocking down Vglut in SLP-AB neurons on flies' preference between fructose and D-glucose, and their corresponding interactions (C: $n$ = 17-21; D: $n$ = 16-20; E: $n$ = 16-18). (F) Effect of knocking down GluClα in AB-FBl8 neurons on flies' preference between fructose and D-glucose, and their corresponding interactions ($n$ = 18-20). (G-H) Knocking down *GluClα* in AB-FBl8 neurons does not affect oscillations in the tip of AB-FBl8 neurons (G: $n$ = 15-19 ; H: $n$ = 17-20). Values represent mean ± SEM. Statistical tests: one-way ANOVA and Tukey post-hoc for behavior and t-test for imaging; ns: $p > 0.05$, ** $p < 0.01$, ***

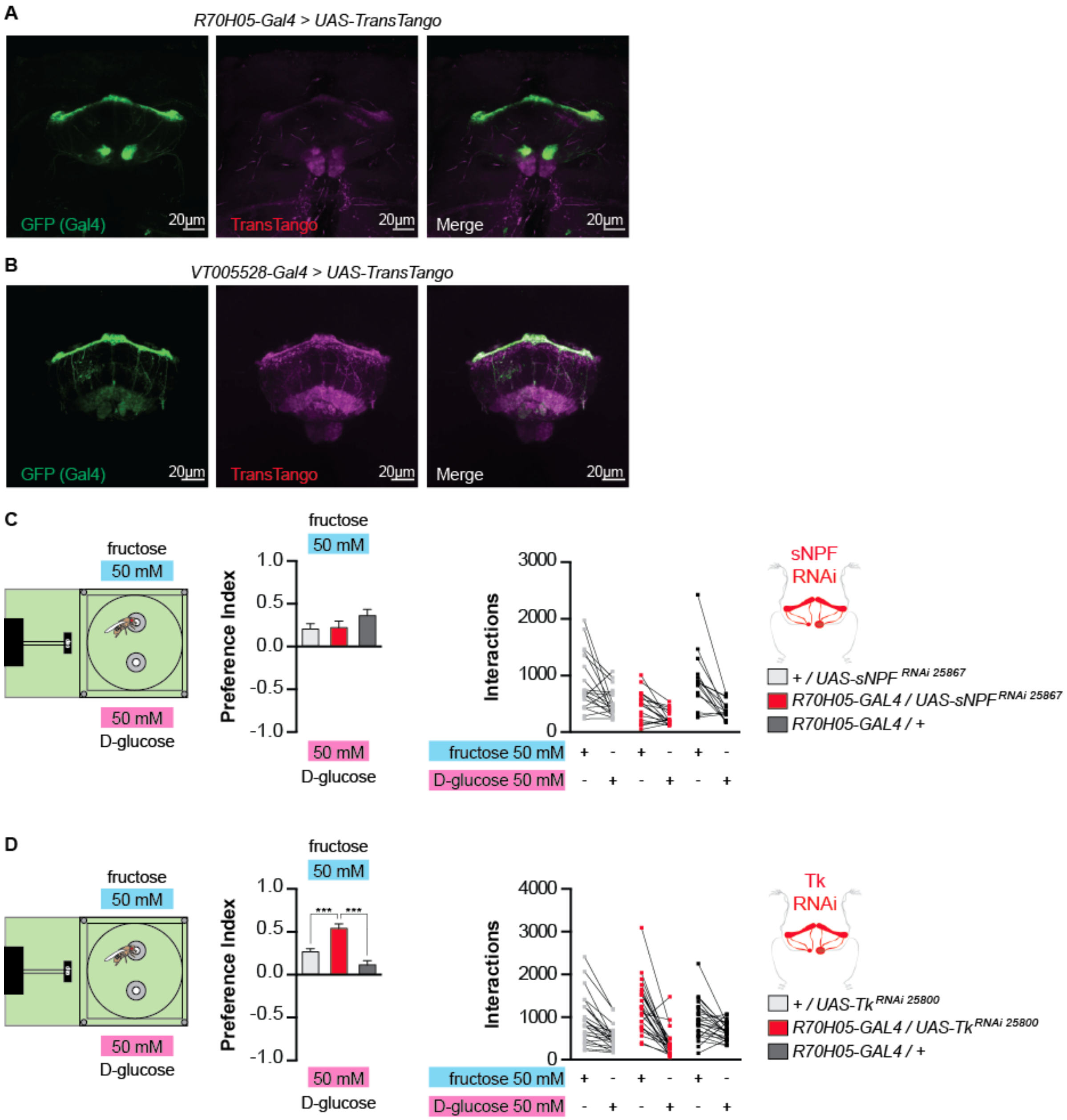


**Figure S7. AB-FBl8 neurons regulate fructose feeding preference through tachykinin secretion, Related to Figure 6**

(A-B) Trans-Tango expression driven by *R70H05-Gal4* (A) and *VT005528-Gal4* (B). (C) Effect of knocking down sNPF in AB-FBl8 neurons on flies' preference between fructose and D-glucose, and their corresponding interactions ($n$ = 15-19). (D) Effect of knocking down tachykinin in AB-FBl8 neurons on flies' preference between fructose and D-glucose, and their corresponding interactions ($n$ = 23-25). Values represent mean±SEM. Statistical tests: one-way ANOVA and Tukey post-hoc; ns: $p > 0.05$, ***$p<0.001$.

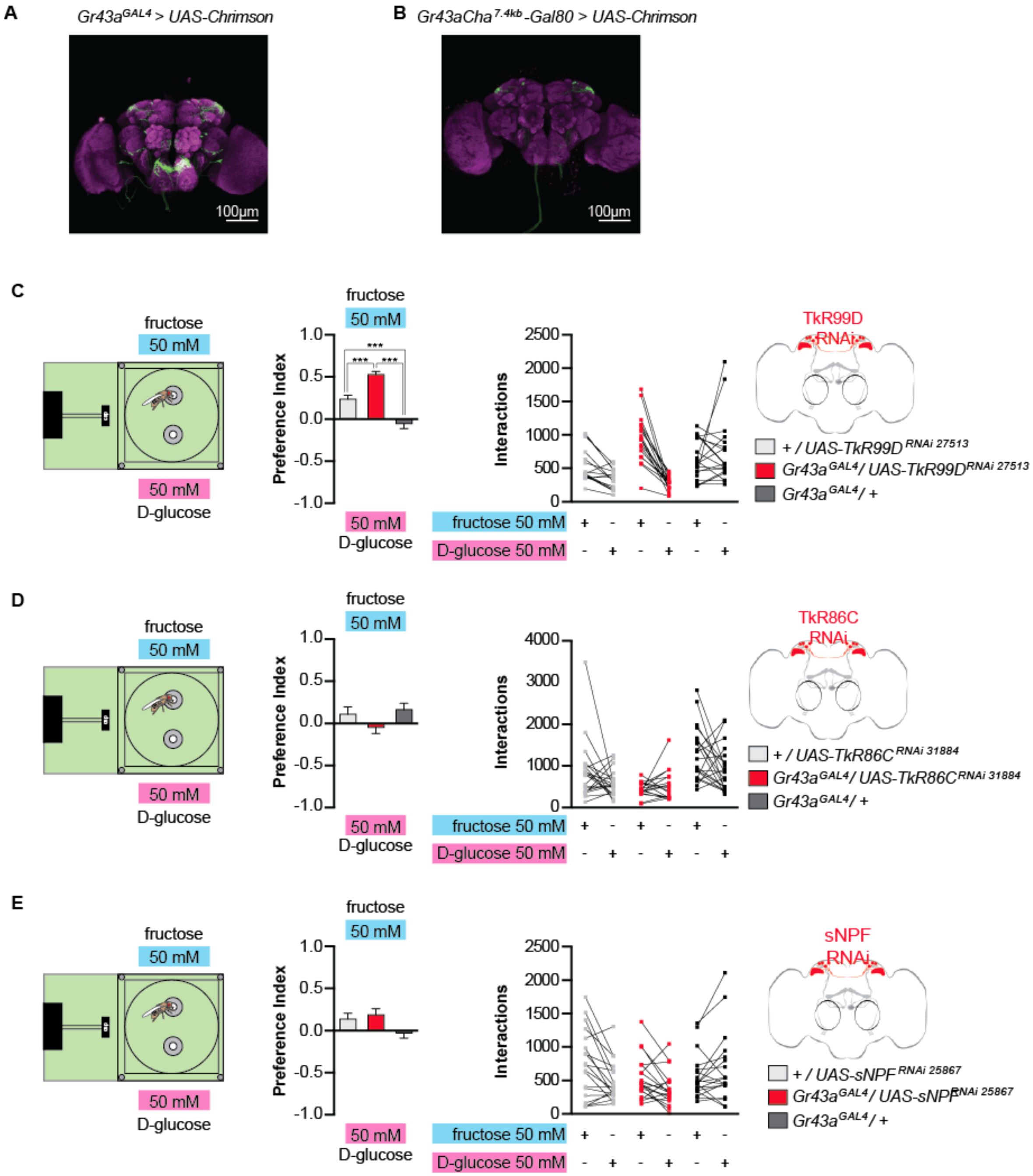


**Figure S8. AB-FBl8 neurons' tachykinin secretion regulates fructose feeding preference through activation of the brain fructose sensor Gr43a, Related to Figure 7**

(A) Immunofluorescent detection of *UAS-Chrimson* driven by *Gr43a$^{GAL4}$*. (B) Immunofluorescent detection of *UAS-Chrimson driven* by *Gr43a-GAL4,Cha$^{7.4kb}$-Gal80*. (C) Effect of knocking down *TkR86C* in Gr43a neurons on flies' preference between fructose and D-glucose, and their corresponding interactions (*n* = 18-21). (D) Effect of knocking down *TkR99D* in Gr43a neurons on flies' preference between fructose and D-glucose, and their corresponding interactions (*n* = 16-19). (E) Effect of knocking down sNPF in Gr43a neurons on flies' preference between fructose and D-glucose, and their corresponding interactions (n = 19-20). Values represent mean±SEM. Statistical tests: one-way ANOVA and Tukey post-hoc; ns: $p > 0.05$, ***$p<0.001$.
